# A new model of endotracheal tube biofilm identifies combinations of matrix-degrading enzymes and antimicrobials able to eradicate biofilms of pathogens that cause ventilator-associated pneumonia

**DOI:** 10.1101/2024.02.20.581163

**Authors:** Dean Walsh, Chris Parmenter, Saskia E Bakker, Trevor Lithgow, Ana Traven, Freya Harrison

## Abstract

Defined as a pneumonia occurring after more than 48 hours of mechanical ventilation via an endotracheal tube, ventilator-associated pneumonia results from biofilm formation on the indwelling tube, seeding the patient’s lower airways with pathogenic microbes such as *Pseudomonas aeruginosa, Klebsiella pneumoniae,* and *Candida albicans.* Currently there is a lack of accurate *in vitro* models of ventilator-associated pneumonia development. This greatly limits our understanding of how the in-host environment alters pathogen physiology and the efficacy of ventilator-associated pneumonia prevention or treatment strategies. Here, we showcase a reproducible model that simulates biofilm formation of these pathogens in a host-mimicking environment, and demonstrate that the biofilm matrix produced differs from that observed in standard laboratory growth medium. In our model, pathogens are grown on endotracheal tube segments in the presence of a novel synthetic ventilator airway mucus (SVAM) medium that simulates the in-host environment. Matrix-degrading enzymes and cryo-SEM were employed to characterise the system in terms of biofilm matrix composition and structure, as compared to standard laboratory growth medium. As seen in patients, the biofilms of ventilator-associated pneumonia pathogens in our model either required very high concentrations of antimicrobials for eradication, or could not be eradicated. However, combining matrix-degrading enzymes with antimicrobials greatly improved biofilm eradication of all pathogens. Our *in vitro* endotracheal tube (IVETT) model informs on fundamental microbiology in the ventilator-associated pneumonia context, and has broad applicability as a screening platform for antibiofilm measures including the use of matrix-degrading enzymes as antimicrobial adjuvants.

**Importance:** The incidence of ventilator-associated pneumonia in mechanically ventilated patients is between 5-40%, increasing to 50-80% in patients suffering from coronavirus disease 2019 (COVID-19). The mortality rate of ventilator-associated pneumonia patients can reach 45%. Treatment of the endotracheal tube biofilms that cause ventilator-associated pneumonia is extremely challenging, with causative organisms able to persist in endotracheal tube biofilm despite appropriate antimicrobial treatment in 56% of ventilator-associated pneumonia patients. Flawed antimicrobial susceptibility testing often means that ventilator-associated pneumonia pathogens are insufficiently treated, resulting in patients experiencing ventilator-associated pneumonia recurrence. Here we present an *in vitro* endotracheal tube biofilm model that recapitulates key aspects of endotracheal tube biofilms, including dense biofilm growth and elevated antimicrobial tolerance. Thus our biofilm model can be used as a ventilated airway simulating environment, aiding the development of anti-ventilator-associated pneumonia therapies and antimicrobial endotracheal tubes that can one day improve the clinical outcomes of mechanically ventilated patients.

## Introduction

Biofilms are congregations of microbial cells coated in a self-produced exopolymeric matrix, consisting of extracellular DNA, exopolysaccharides, matrix proteins, and lipids; the biofilm aggregate is often surface-attached, but can be free floating (1). The microbes ensconced in a biofilm are extremely difficult to treat with antimicrobial therapy due to: i) antimicrobial diffusion being impeded by the biofilm matrix (2–6), ii) an increased population of persister cells within biofilms (7, 8), iii) hypermutability and horizontal gene transfer allow for the acquisition of genetic mechanisms of antimicrobial resistance (8–10).

Medical-device related infections account for a quarter of all nosocomial infections (11). In the case of critically ill patients intubated with endotracheal tubing to facilitate mechanical ventilation of the airways, intubation increases the risk of pulmonary infection up to 20-fold (12, 13). Intubation compromises the local immune response, prevents the cough reflex, impedes mucociliary clearance of entrapped microorganisms, and damages the tracheal epithelium. Following intubation, oropharyngeal flora and nosocomial pathogens colonise the trachea and form biofilms on the endotracheal tubing. Furthermore, accumulation of contaminated secretions above the endotracheal tube cuff leads to microaspiration of these secretions, seeding biofilm to other regions of the endotracheal tube and airways, potentially resulting in ventilator-associated pneumonia (also called VAP) (14–16).

Many oral commensal and hospital-acquired pathogen species can form monospecies or polymicrobial biofilms on endotracheal tubes *in vivo* (17, 18). A previous study found that the pathogens found in the lung were identical to those constituting endotracheal tube biofilms in 70% of ventilator-associated pneumonia patients (19). *Staphylococcus aureus*, *Klebsiella pneumoniae*, *Acinetobacter baumannii*, and *Pseudomonas aeruginosa* are among the pathogens most commonly associated with ventilator-associated pneumonia, particularly with late-onset disease in which multidrug resistant strains become prevalent (20–22). *Candida albicans*, an oral commensal and opportunistic pathogen, rarely causes ventilator-associated pneumonia but frequently colonises endotracheal tubes (17, 20, 23).

Eradication of endotracheal tube biofilms is extremely challenging, with one study finding causative organisms persisting on endotracheal tubes despite appropriate antimicrobial treatment in 56% of ventilator-associated pneumonia patients (23). Furthermore, ventilator-associated pneumonia recurrence occurs in 26% of patients, with on average 2-5 recurrences per patient; this is often due to failed treatment of the initial pathogen (24). While antimicrobial coated endotracheal tubes can prevent the onset of ventilator-associated pneumonia, few of these have received FDA approval (25). Whilst coated endotracheal tubes reduce ventilator-associated pneumonia incidence, there is little evidence to suggest they reduce crucial patient outcomes including hospital stay, mechanical ventilation duration, or patient mortality. This makes it difficult for clinicians to justify the higher costs of coated endotracheal tubes (26), demonstrating the need for further innovation in endotracheal tube development.

Relapses in infection can often be attributed to flawed antimicrobial susceptibility testing (27–30). Conventional antimicrobial susceptibility testing methods cannot always predict the efficacy of antibiotic treatment *in vivo* (27), and do not factor in biofilm or polymicrobial communities (28, 30). To bridge this disconnect between conventional antimicrobial susceptibility testing and the *in vivo* environment, researchers have developed biofilm models that attempt to capture the infection environment and increase diagnostic accuracy. These include Calgary devices in which bacteria can form biofilms on plastic pegs (31, 32), however these methods do not provide much improvement on standard antimicrobial susceptibility testing (33). More recent biofilm models, such as our *ex vivo* model of the cystic fibrosis airways, which combines pig lung tissue with synthetic cystic fibrosis mucus, produce biofilms with gene expression profiles that resemble those observed in patient sputum (2, 34–43). The *in vitro* and *ex vivo* models developed to better mimic the airways of individuals with cystic fibrosis (44) stands in stark contrast to ventilator-associated pneumonia, where animal models are relied on for the study of the disease (45, 46).

Here, we showcase a novel *in vitro* biofilm model featuring newly developed synthetic ventilated airway mucus growth medium (SVAM) and serum-coated endotracheal tubes (Fig 1). This model simulates the nutritional and material environment that pathogens are exposed to as they colonise patient airways. We found that biofilms of *P. aeruginosa*, *K. pneumoniae* and *C. albicans* cultured in this model were much harder to eradicate with antimicrobial treatment compared to culture in standard antimicrobial susceptibility testing conditions. Furthermore, cryo-SEM and enzymatic degradation assays revealed that biofilm structure and matrix composition differed depending on the growth environment. We then used our model to screen combinations of antibiotics and matrix-degrading enzymes against biofilms of ventilator-associated pneumonia pathogens. This combination treatment resulted in improved biofilm eradication. Collectively, this study illustrates how a tailored model of endotracheal tube biofilm may be used to test emerging therapies designed to prevent biofilm formation and the onset of ventilator-associated pneumonia in mechanically ventilated patients.

**Figure 1:**
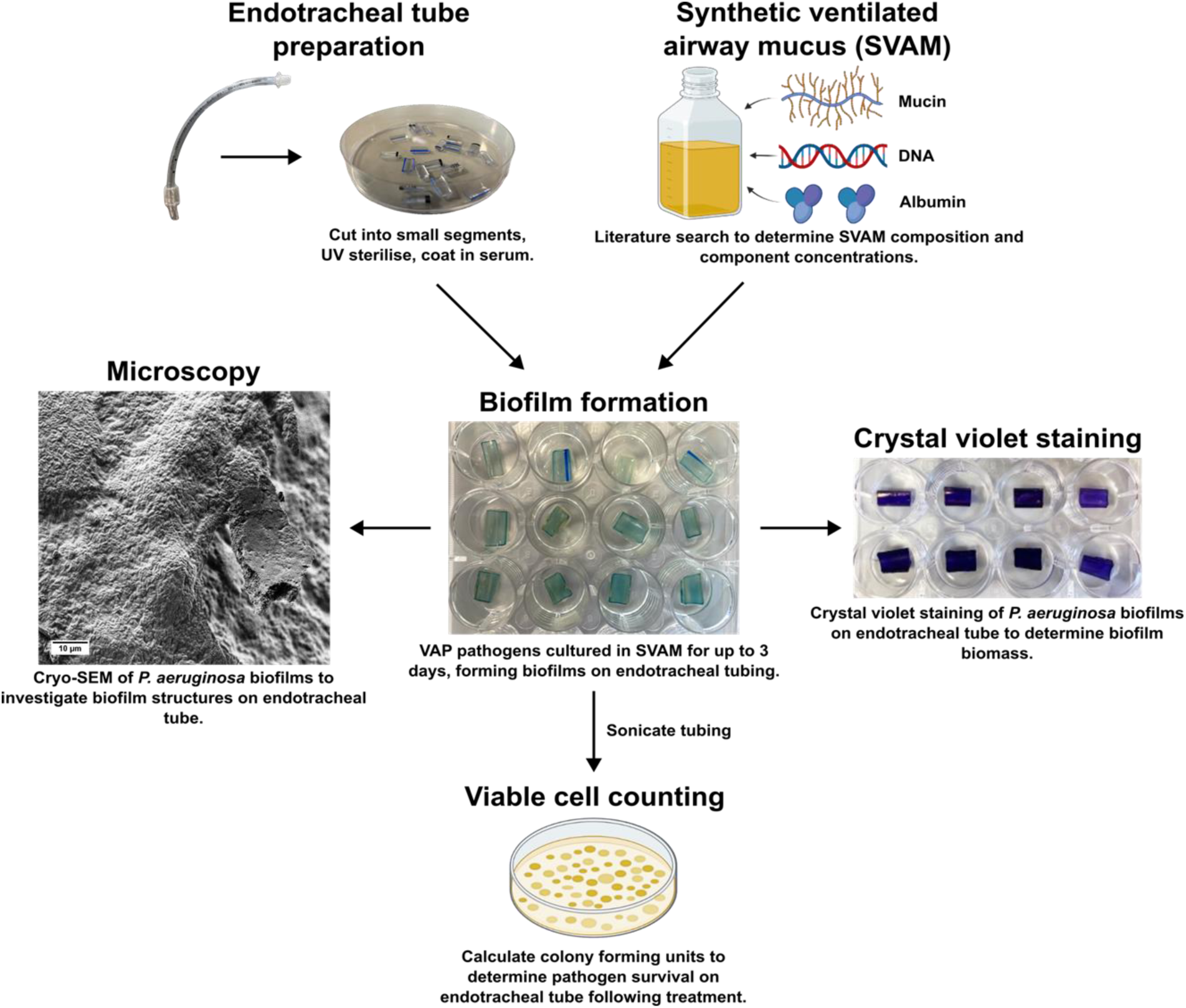
Schematic and applications of the *in vitro* endotracheal tube biofilm model. *P. aeruginosa* biofilms are shown as an example to illustrate the model. Figure created with BioRender.com.

## Materials and Methods

### Strains and culture conditions

Cultures of *P. aeruginosa* PA14 and *K. pneumoniae* B5055 were grown aerobically at 37°C overnight on lysogeny broth (LB) agar plates for solid culture, and overnight in LB broth at 37°C with shaking at 150 rpm for liquid culture. *C. albicans* SC5314 solid cultures were grown aerobically overnight on yeast extract peptone dextrose (YPD) agar plates at 37°C, whilst liquid cultures were grown overnight in YPD broth at 37°C with shaking at 150 rpm.

### Synthetic ventilated airway mucus (SVAM) medium

Individual stocks of 9.6 mg/mL salmon sperm DNA, 17 mM Fe(II)SO_4_·7H_2_O, 95 mM ZnSO_4_·7H_2_O, 9.5 mM CuSO_4_·5H_2_O, 4 mg/mL sialic acid, and 100 mg/mL human lysozyme were filter sterilised (0.22 µm, PES filter). A 20 mg/mL stock of type III porcine gastric mucin (PGM) was prepared and sterilised by autoclaving. Human lysozyme stock was stored at −20°C, all other stocks were stored at 4°C. Preparation of the SVAM base stock is detailed in Table 1. Preparation of the completed SVAM medium, which is prepared fresh in aseptic conditions immediately prior to each experiment, is detailed in Table 2. Final concentrations of components in stocks and in the final SVAM medium is shown in Table S1. Further details and suppliers of reagents are shown in Table S2.

**Table 1:** Components and method for preparing the SVAM base stock.

| Component | Amount required 500 mL SVAM base stock | Protocol |
| --- | --- | --- |
| CaCl<sub>2</sub> | 0.236 g | Add each component individually in the order listed to 480 mL of dH<sub>2</sub>O, ensuring each component is |
| Bovine serum albumin | 10.5 g |  |
| NaCl | 3.33 g |  |
| Na<sub>2</sub>HPO<sub>4</sub> | 0.86 g |  |
| NaH<sub>2</sub>PO<sub>4</sub> | 0.206 g |  |
| **K<sub>2</sub>SO<sub>4</sub>** | 1.087 g | fully dissolved before adding the next component.<br>• NaOH to pH 6.7-6.8.<br>• Add dH<sub>2</sub>O to give a final volume of 500 mL.<br>• Filter sterilise (0.22 µm, PES filter) and store at 4°C. |
| **KCl** | 0.131 g |  |
| **Casamino acids** | 7.5 g |  |
| **MOPS** | 3.139 g |  |
| **MgCl<sub>2</sub>·6H<sub>2</sub>O** | 0.725 g |  |
| **Glucose** | 1.08 g |  |
| **DPTA** | 0.009 g |  |
| **GlcNAc** | 5.25 g |  |
| **Fe(II)SO<sub>4</sub>·7H<sub>2</sub>O** | 1.5 mL of 17 mM stock |  |
| **ZnSO<sub>4</sub>·7H<sub>2</sub>O** | 1.5 mL of 95 mM stock |  |
| **CuSO<sub>4</sub>·5H<sub>2</sub>O** | 1.5 mL of 9.5 mM stock |  |
| **Sialic acid** | 1.5 mL of 4 mg/mL stock |  |

**Table 2:** Preparation of the final SVAM medium, prepared fresh for each experiment.

| **Component** | **Amount required for 30 mL of completed SVAM medium** | **Comments** |
| --- | --- | --- |
| **SVAM base stock (filter sterile)** | 10 mL | SVAM base stock always makes up 33.3% of final volume. |
| **9.6 mg/mL salmon sperm DNA stock (filter sterile)** | 3 mL | DNA stock always makes up 10% of final volume |
| **20 mg/mL porcine stomach mucin (type III) stock (autoclaved)** | 15 mL | Mucin stock always makes up 50% of final volume. |
| **100 mg/mL lysozyme stock (filter sterile)** | 15 µL | Add lysozyme to a final concentration of 48 µg/mL. |
| **NaOH** | Add until final pH of 6.7-6.8 is reached. | Aseptically check pH. |
| **dH<sub>2</sub>O** | Add until 30 mL final volume achieved. |  |

### Antimicrobial susceptibility testing using standard platforms

Cation-adjusted Mueller Hinton broth (caMHB) (Sigma-Aldrich) or SVAM was inoculated with *P. aeruginosa*, *K. pneumoniae*, or *C. albicans* to a final OD_600nm_ of 0.05 into 96 well plates (Corning, 3596). *P. aeruginosa* was tested against gentamicin, *K. pneumoniae* against colistin, and *C. albicans* against amphotericin B. All organisms were grown aerobically throughout standard antimicrobial susceptibility testing. All antimicrobials were tested at the range 0.125–256 µg/mL. For planktonic cultures, minimum inhibitory concentrations (MICs) were determined by broth microdilution in accordance with EUCAST guidelines (European Committee on Antimicrobial Susceptibility Testing: www.eucast.org). For biofilms, following the same inoculation procedure described above, minimum biofilm eradication concentrations (MBEC) were determined using a peg lid Calgary biofilm device (peg lid: Nunc (445497), plate: Nunc (269787)) as previously described (31).

### Antimicrobial susceptibility testing in the in vitro endotracheal tube biofilm model

Under aseptic conditions, endotracheal tubes (siliconized PVC, cuffed, 8 mm, IMS Euro) were cut into 1 cm rings. Each ring was then cut into 6 equal segments. Segments were sterilised under short-wave UV light (Carlton Germicidal Cabinet) for 10 minutes. Fetal bovine serum (FBS) was aseptically poured over the endotracheal tube segments until immersed, and the segments were sealed in a petri dish and left overnight at 4°C to become coated in serum proteins.

Serum-coated endotracheal tube segments were added to the wells of 24 well plates (Corning Costar, CLS3527). Endotracheal tube segments were immersed in 0.5 mL of SVAM medium, inoculated with 0.05 OD_600_ of relevant microorganisms. The plates were then incubated at 37°C, 5% CO_2_ for 48 hours to allow biofilm formation. Following incubation, biofilm coated endotracheal tube segments were transferred to fresh SVAM medium containing relevant antimicrobials. This transfer simulates the periodic removal of secretions from the airways of ventilated patients and the build-up of fresh secretions. Antimicrobial exposed biofilms were incubated for a further 24 hours at 37°C and 5% CO_2_. Biofilms were transferred to fresh plates, washed with 1 mL of PBS, sonicated at 50 Hz (Grant XUBA1 sonicating water bath) for 15 minutes and then scraped with sterile pipette tips to remove biofilm. Suspensions were plated onto relevant growth medium and colony forming units/mL (CFU/mL) were calculated to determine the viability of endotracheal tube biofilm.

### Cryo-scanning electron microscopy (cryo-SEM) of endotracheal tube biofilms

*P. aeruginosa*, *K. pneumoniae*, and *C. albicans* were grown on serum-coated endotracheal tubes in either SVAM and LB (*P. aeruginosa* and *K. pneumoniae*), or SVAM and YPD (*C. albicans*), for 72 hours at 37°C, 5% CO_2_. Biofilms were fixed for 2 hours in 2% paraformaldehyde in PBS. Imaging was carried out using a Zeiss Crossbeam 550 FIB-SEM equipped with a Quorum 3010 cryo sample preparation system. Following cryo preparation in slushy nitrogen, biofilm samples were sublimated for 5 minutes at −90°C and coated with platinum for 1 minute at a current of 10 mA. Endotracheal tube biofilm samples were tilted at 30°, the accelerating voltage of the SEM was maintained at 2 kV, and working distance maintained at approximately 4-5 mm for each image.

### Susceptibility of endotracheal tube biofilms to matrix-degrading enzymes

Serum-coated endotracheal tube segments were added to the wells of 24 well plates and immersed in 0.5 mL of LB (*P. aeruginosa, K. pneumoniae*), YPD (*C. albicans*), or SVAM, inoculated with 0.05 OD_600_ of relevant microorganisms. Plates were incubated at 37°C, 5% CO_2_ for 48 hours to allow biofilm formation. Following incubation, biofilm coated endotracheal tubes were transferred to fresh growth medium containing either 100 µg/mL DNase I, 1 mg/mL proteinase K, or 10% (w/v) glycoside hydrolases (5% (w/v) cellulase, 5% (w/v) α-amylase). Further details on enzyme activity and suppliers found in Table S2. Biofilms were then incubated for a further 24 hours at 37°C and 5% CO_2_.

Endotracheal tube segments were removed from wells and immersed in 0.05% (v/v) crystal violet solution for 15 minutes, washed in PBS and left to dry for 30 minutes in a laminar flow cabinet. Endotracheal tubes were then immersed in 30% acetic acid for 15 minutes to solubilise crystal violet. Biomass was determined by reading absorbance at 550 nm.

### Testing antimicrobial and matrix-degrading enzyme combinations in the in vitro endotracheal tube biofilm model

Serum-coated endotracheal tube segments were added to the wells of 24 well plates and immersed in 0.5 mL of SVAM medium, inoculated with 0.05 OD_600_ of relevant microorganism. Plates were incubated at 37°C, 5% CO_2_ for 48 hours to allow biofilm formation. Following incubation, biofilm coated endotracheal tubes were transferred to fresh SVAM medium containing relevant antimicrobials. Relevant wells were also supplemented with enzymes as described above. Biofilms were incubated for a further 24 hours at 37°C and 5% CO_2_. Endotracheal tubes were transferred to fresh plates, washed with 1 mL of PBS, sonicated for 15 minutes and scraped with sterile pipette tips to remove biofilm. Following removal of endotracheal tubes from treatment wells and into wells of PBS, CFU/mL was determined. CFU/mL from the endotracheal tubes determine the viability of biofilm still attached to the endotracheal tube. Total CFU/mL were calculated from the sum of the dispersed and biofilm CFUs.

### Data analysis

Raw data are provided in the Data Supplement (will be supplied upon article acceptance). Data presentation and statistical analyses were conducted in GraphPad Prism 10. Two-way ANOVAs with Dunnett’s multiple comparisons tests were carried out for biofilm dispersal experiments, with P < 0.05 considered significant.

## Results

### Synthetic ventilated airway mucus medium (SVAM) increases the MIC and MBEC of clinically relevant antibiotics against ventilator-associated pneumonia pathogens

Gentamicin and colistin were chosen to treat *P. aeruginosa* and *K. pneumoniae*, respectively, as both antibiotics are frequently used to treat multidrug resistant Gram-negative bacteria (47, 48) and carbapenem resistant ventilator-associated pneumonia infections (47, 49). Amphotericin B was used for *C. albicans* treatment due to its use in airway decolonisation of *C. albicans* in mechanically ventilated patients, which is known to reduce ventilator-associated pneumonia incidence and improving prognosis (50).

Although not as well characterised as cystic fibrosis sputum, existing literature does suggest that mucus in the ventilated airways is a distinct growth environment that necessitates a bespoke growth medium. These distinctions include much higher mucin and glucose concentrations compared to those in cystic fibrosis sputum (51–54). The SVAM medium recipe was developed following an extensive review of existing literature (55). The effect of each SVAM component on the growth and biofilm formation of *P. aeruginosa*, *K. pneumoniae* and *C. albicans* is shown in Figures S1-S3, following a protocol detailed in the Supplementary Methods.

In a standard broth microdilution assay, culture medium used affected the concentration of antimicrobial required to inhibit growth of all species investigated. In caMHB, all species were more susceptible to their respective antimicrobial treatments than when grown in SVAM (Table 3). When grown in SVAM medium, the gentamicin MIC of *P. aeruginosa* was 4-fold higher than in caMHB (caMHB: 4 µg/mL, SVAM: 16 µg/mL). Both *K. pneumoniae* and *C. albicans* displayed an 8-fold increase in MICs to colistin (caMHB: 1 µg/mL, SVAM: 8 µg/mL) and amphotericin B (caMHB: 0.25 µg/mL, SVAM: 2 µg/mL), respectively.

**Table 3:** Minimum inhibitory concentrations of clinically relevant antimicrobials against ventilator-associated pneumonia pathogens.

| Organism | Antimicrobial | Broth microdilution (caMHB) MIC ($\mu\text{g/mL}$) | Broth microdilution (SVAM) MIC ($\mu\text{g/mL}$) |
| --- | --- | --- | --- |
| *P. aeruginosa* PA14 | Gentamicin | 4 | 16 |
| *K. pneumoniae* B5055 | Colistin | 1 | 8 |
| *C. albicans* SC5314 | Amphotericin B | 0.25 | 2 |

As expected, the antimicrobial concentrations required to eradicate biofilms grown in a peg lid biofilm Calgary device exceeded the MIC required in planktonic culture. MBECs for all pathogens were higher in SVAM than in caMHB: for *P. aeruginosa* (Fig 2A, 2D) and *K. pneumoniae* (Fig 2B, 2E), the respective antibiotics were half as effective (*P. aeruginosa*: 32 µg/mL to 64 µg/mL gentamicin, *K. pneumoniae*: 128 µg/mL to 256 µg/mL colistin) and for *C. albicans* amphotericin B was 4-fold less effective (from 8 µg/mL in caMHB, to 32 µg/mL in SVAM).

**Figure 2:**
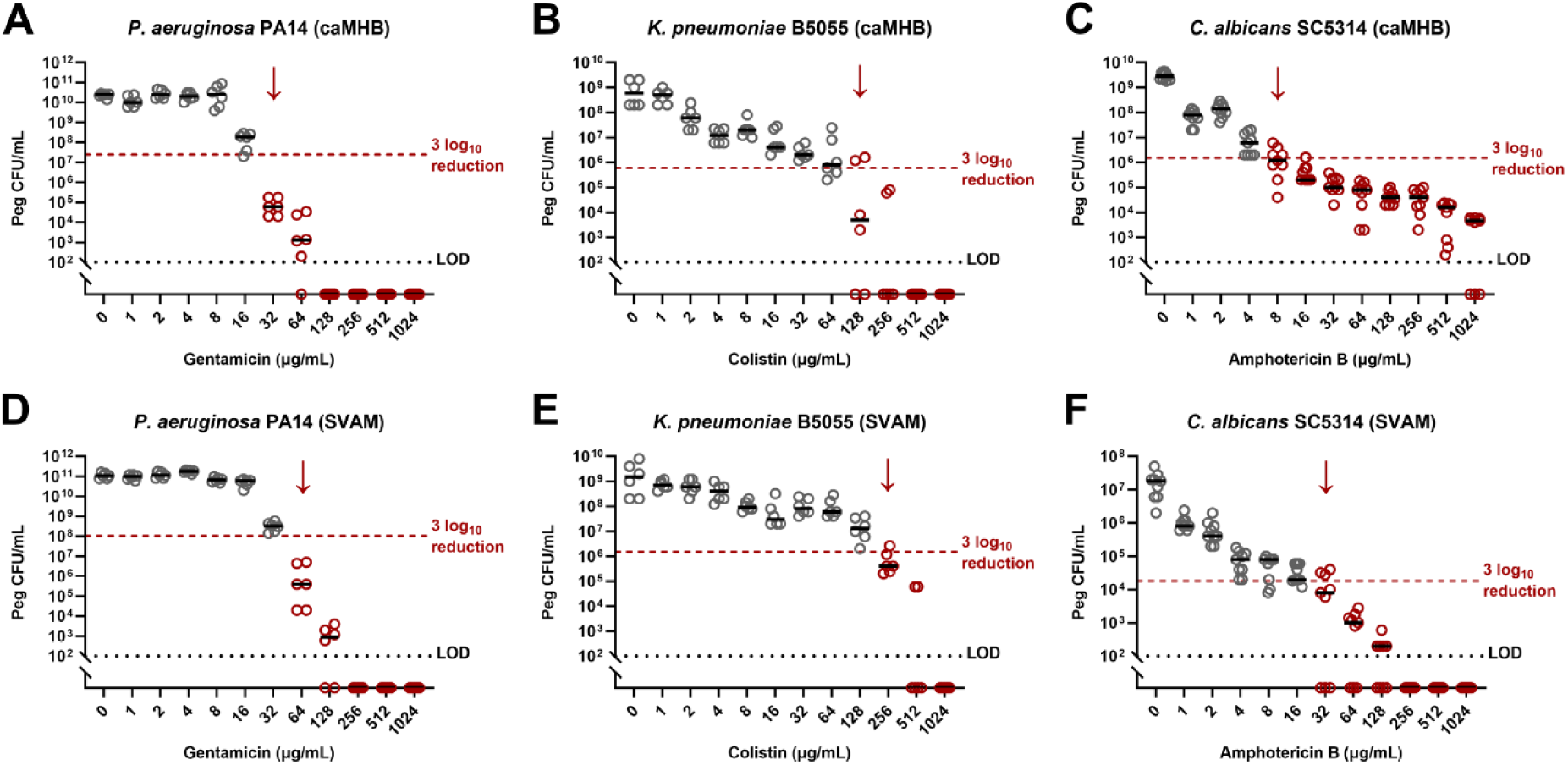
The effect of growth medium on the antimicrobial susceptibility of biofilms grown in a Calgary biofilm device. Biofilms of ventilator-associated pneumonia pathogens were established in peg lid Calgary biofilm devices, in either caMHB or SVAM, for 48 hours. Peg lids were then transferred to fresh medium with antimicrobials for a further 24 hours. Biofilms were removed from pegs and CFUs counted to determine endpoint viability. The minimum biofilm eradication concentration is defined as the antimicrobial concentration needed to achieve an average 3 log10 reduction in CFUs relative to untreated controls. **A)** Endpoint CFUs of *P. aeruginosa* biofilms grown in caMHB. **B)** Endpoint CFUs of *K. pneumoniae* biofilms grown in caMHB. **C)** Endpoint CFUs of *C. albicans* biofilms grown in caMHB. **D)** Endpoint CFUs of *P. aeruginosa* biofilms grown in SVAM. **E)** Endpoint CFUs of *K. pneumoniae* biofilms grown in SVAM. **F)** Endpoint CFUs of *C. albicans* biofilms grown in SVAM. Dotted line denotes the 3 log10 reduction threshold. Black dotted line denotes the limit of detection (LOD). Red arrow highlights the minimum biofilm eradication concentration, n = 3 biological repeats (experiment repeated on different days with fresh starting cultures), with 2 technical repeats (*P. aeruginosa* and *K. pneumoniae*) or 3 technical repeats (*C. albicans*) for each biological replicate.

Further increases in antimicrobial tolerance were observed when pathogens formed biofilms on serum-coated endotracheal tubes in SVAM. Relative to the peg lid biofilms, eradication of *P. aeruginosa* (Fig 3A) and *C. albicans* (Fig 3C) required higher concentrations of gentamicin (4x higher; 256 µg/mL), and amphotericin B (8x higher; 512 µg/mL), respectively. However, while the presence of serum-coated endotracheal tubes increased the antimicrobial tolerance of some ventilator-associated pneumonia pathogens, this was not universal, with *K. pneumoniae* displaying no change in colistin susceptibility (256 µg/mL) between the peg lid Calgary device (Fig 2E) and the endotracheal tube model (Fig 3B).

**Figure 3:**
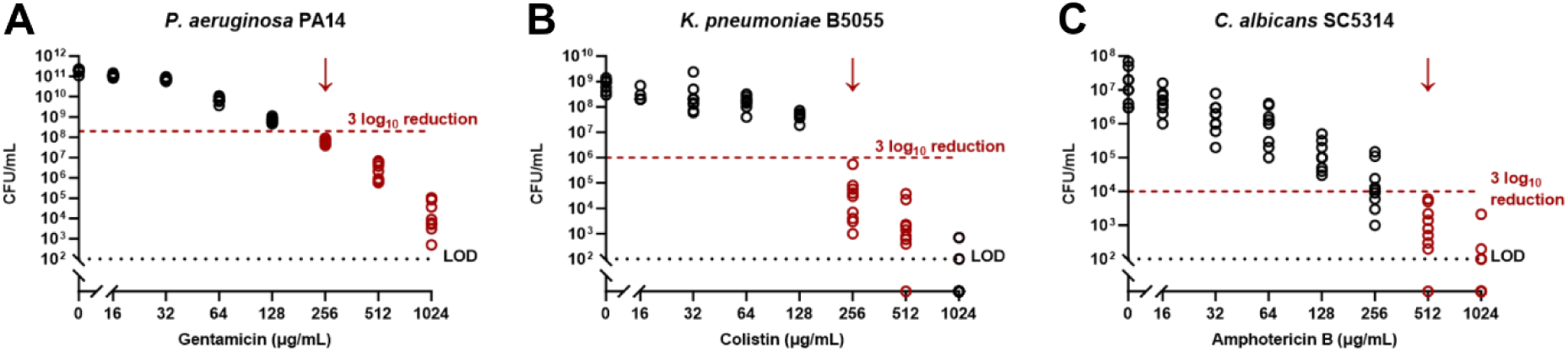
Susceptibility of biofilms to clinically relevant antimicrobials in the *in vitro* endotracheal tube model. Ventilator-associated pneumonia pathogens were grown on segments of serum-coated endotracheal tube in SVAM for 48 hours to establish biofilms. Endotracheal tubes were then transferred to fresh SVAM with antimicrobials for a further 24 hours. Biofilms were removed from endotracheal tube segments and CFUs counted to determine endpoint viability. MBECs were defined as the first antimicrobial concentration to achieve an average 3 log10 reduction in CFUs relative to untreated controls. **A)** Endpoint CFUs of *P. aeruginosa* biofilms. **B)** Endpoint CFUs of *K. pneumoniae* biofilms. **C)** Endpoint CFUs of *C. albicans* biofilms. Red dotted line denotes the 3 log10 reduction threshold. Black dotted line denotes the limit of detection (LOD). Red arrow highlights the minimum biofilm eradication concentration, n = 3 biological repeats, with 3 technical repeats for each biological replicate.

### SVAM impacts the structure, matrix composition and susceptibility to matrix degrading enzymes of biofilms grown on endotracheal tubes

Since biofilms grown in the *in vitro* endotracheal tube model were more tolerant to antimicrobial therapy, we sought to discern whether changes in biofilm structure and matrix composition might explain this. Biofilms of ventilator-associated pneumonia pathogens were grown on endotracheal tubes either in SVAM to mimic the ventilator-associated pneumonia environment, or in standard laboratory growth media (LB for *P. aeruginosa* and *K. pneumoniae*; YPD for *C. albicans*) and Cryo-SEM was employed to visualise biofilm ultrastructure.

Cryo-SEM of *P. aeruginosa* biofilms showed that dense biofilm coats the endotracheal tube in either growth medium. However, *P. aeruginosa* endotracheal tube biofilms grown in LB appeared flat and homogenous, whereas those grown in SVAM had a varied topography, with microcolony structures covering the tube (Fig 4A). When *K. pneumoniae* biofilms were grown in LB, only small aggregates of biofilm and single cells could be found on the endotracheal tube (Fig 4B). Conversely, SVAM resulted in *K. pneumoniae* biofilms covered in a thick matrix. Furthermore, *K. pneumoniae* cells in LB-grown biofilms had a smooth surface, whereas those grown in SVAM had a rough cell surface, which is likely due to a thicker coating of capsular polysaccharide. Lastly, YPD-grown endotracheal tube biofilms of *C. albicans* were largely composed of layers of hyphae (Fig 4C), whereas those grown in SVAM instead favoured yeast cells and pseudohyphae. This was particularly apparent in the images of freeze-fractured *C. albicans* biofilms, showing a biofilm interior composed of many layers of yeast cells and pseudohyphae. *C. albicans* biofilms grown in SVAM were also much thicker than those grown in YPD. All imaged biofilms were grown on serum-coated ETTs, indicating that structural differences were solely due to differences in growth medium, rather than the presence of serum.

**Figure 4:**
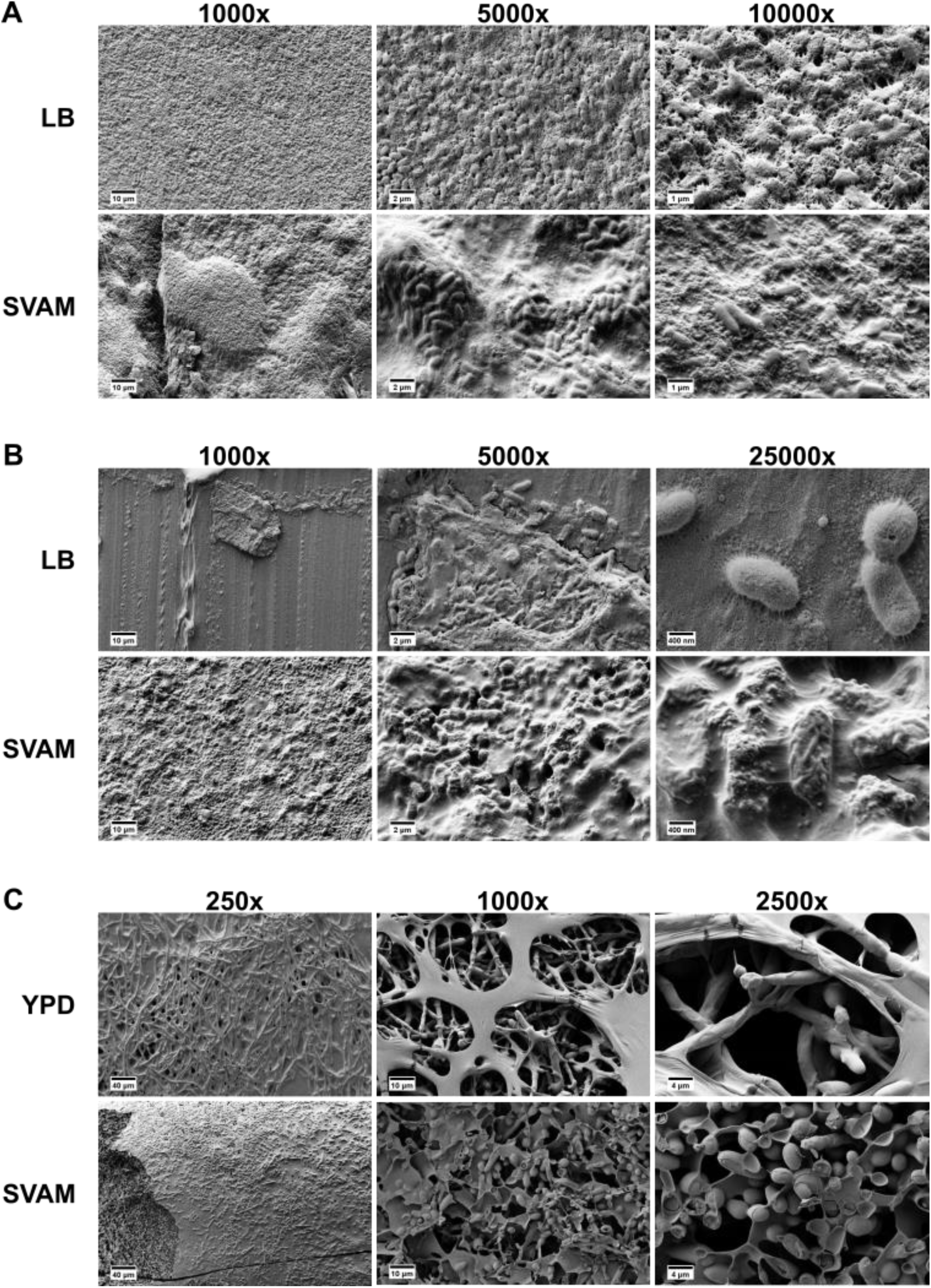
Cryo-SEM shows that the endotracheal tube environment alters the biofilm structure of *P. aeruginosa*, *K. pneumoniae*, and *C. albicans*. **A)** Representative images of *P. aeruginosa* biofilms grown on serum-coated endotracheal tube either in LB or SVAM. **B)** Representative images of *K. pneumoniae* biofilms grown on serum-coated endotracheal tube either in LB or SVAM. **C)** Representative images of *C. albicans* biofilms grown on serum-coated endotracheal tube either in YPD or SVAM. Relevant magnifications are shown above each set of images.

Matrix composition was determined by treating established endotracheal tube biofilms with hydrolytic enzymes. DNase I, proteinase K, and glycoside hydrolases degrade extracellular DNA, matrix proteins, and exopolysaccharides, respectively. Following enzymatic treatment, crystal violet staining was used to detect residual biofilm. Treatment with glycoside hydrolases was able to significantly disperse *P. aeruginosa* biofilms grown in either LB or SVAM from the endotracheal tube segment (Fig 5A). Conversely, *K. pneumoniae* (Fig 5B) and *C. albicans* (Fig 5C) biofilms demonstrated differences in matrix degrading enzyme susceptibilities when grown in different media. Biofilms of *K. pneumoniae* grown in LB were sensitive to dispersal by glycoside hydrolases, whereas SVAM-grown *K. pneumoniae* biofilms were impervious to dispersal by glycoside hydrolases. In the case of *C. albicans*, growth in YPD established a biofilm sensitive to DNase but insensitive to glycoside hydrolases, whereas biofilms established in SVAM were glycoside hydrolase sensitive but recalcitrant to DNase treatment. Susceptibility of *K. pneumoniae* biofilms, *P. aeruginosa* biofilms and *C. albicans* to proteinase K dispersal was similar irrespective of the growth medium used to establish the biofilm. Thus, the macromolecular composition of the extracellular matrix differs in biofilms established in SVAM.

**Figure 5:**
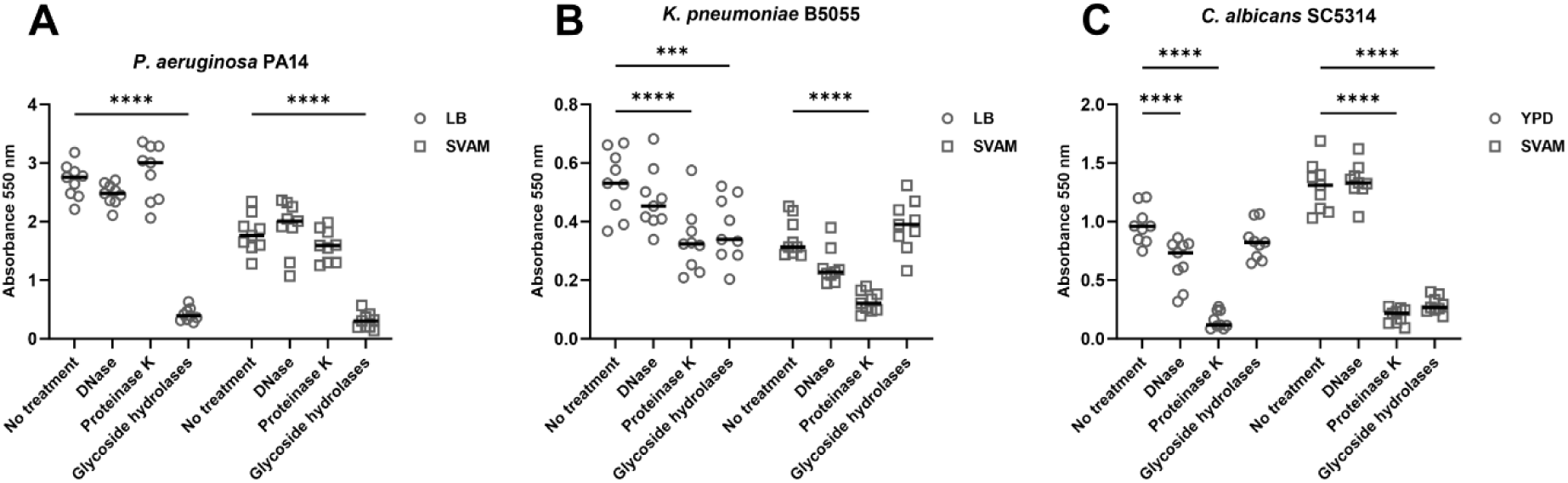
Susceptibility of endotracheal tube biofilms to matrix degrading enzymes when grown in standard laboratory medium vs SVAM medium. Ventilator-associated pneumonia pathogens were grown on segments of serum-coated endotracheal tube in either LB (*P. aeruginosa* and *K. pneumoniae*), YPD (*C. albicans*), or SVAM for 48 hours to establish biofilms. Endotracheal tubes were then transferred to fresh medium and treated with either 100 µg/mL DNase, 1 mg/mL proteinase K, or 10% glycoside hydrolases for a further 24 hours. Endotracheal tube biofilms were then stained with crystal violet and absorbance measured at 550 nm to determine changes in biofilm biomass. **A)** Biomass of *P. aeruginosa* endotracheal tube biofilms (enzyme: F3,64 = 164.6, P < 0.0001; medium: F1,64 = 94.85, P < 0.0001; interaction F3,64 = 11.27, P < 0.0001). **B)** Biomass of *K. pneumoniae* endotracheal tube biofilms (enzyme: F3,64 = 17.46, P < 0.0001; medium: F1,64 = 52.29, P < 0.0001; interaction F3,64 = 7.24, P = 0.0003). **C)** Biomass of *C. albicans* endotracheal tube biofilms (enzyme: F3,64 = 159.9, P < 0.0001; medium: F1,64 = 13.98, P = 0.0004; interaction F3,64 = 56.58, P < 0.0001). Statistical significance was determined by a two-way ANOVA with Dunnett’s multiple comparisons test. **** denotes P ≤ 0.0001, *** denotes P ≤ 0.001, n = 3 biological repeats, with 3 technical repeats for each biological replicate.

### Combining antimicrobials with matrix degrading enzymes can increase eradication of biofilms grown in the in vitro endotracheal tube model

We sought to evaluate combined treatments with matrix degrading enzymes and antimicrobial drugs to overcome the antimicrobial tolerance in the endotracheal tube biofilms. There is clinical potential for such approaches, with recombinant human DNase (rhDNase) already in use as a mucolytic therapy in the airways of cystic fibrosis patients (56, 57). Thus, we investigated the efficacy of DNase in conjunction with gentamicin to treat *P. aeruginosa* biofilms. While DNase had little impact alone in enzymatic dispersal experiments (Fig 5A), its presence alongside gentamicin had a synergistic effect on eradication of *P. aeruginosa*: DNase plus 128 µg/mL gentamicin was better at reducing the total viable *P. aeruginosa* population than either 128 µg/mL or 256 µg/mL gentamicin alone (Fig 6C). Importantly, this treatment killed cells in both the dispersed (Fig 6A) and biofilm (Fig 6B) populations of *P. aeruginosa*, showing that this combination did not simply disperse the biofilm, it also improved killing by gentamicin.

**Figure 6:**
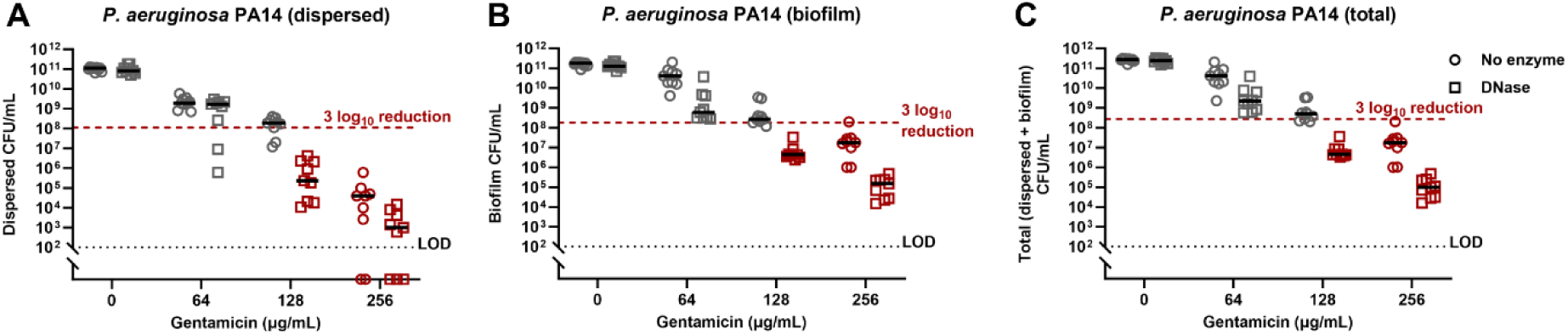
Treating *P. aeruginosa* grown in the *in vitro* endotracheal tube biofilm model with a combination of DNase and gentamicin. Biofilms of *P. aeruginosa* were established on segments of serum-coated endotracheal tube in SVAM for 48 hours. Endotracheal tubes were then transferred to fresh medium and treated with either 100 µg/mL DNase and differing gentamicin concentrations for a further 24 hours. Endpoint CFUs of the dispersed bacteria in the surrounding medium **(A)**, the endotracheal tube associated biofilm **(B)**, and the total population (dispersed + biofilm) **(C)** were acquired. Red dotted line denotes the 3 log10 reduction threshold. Black dotted line denotes the limit of detection (LOD), n = 3 biological repeats, with 3 technical repeats for each biological replicate.

Glycoside hydrolases significantly dispersed both *P. aeruginosa* and *C. albicans* endotracheal tube biofilms grown in SVAM (Fig 5A, 5C). Therefore, we evaluated the effect of combining glycoside hydrolases with gentamicin (*P. aeruginosa*) or amphotericin B (*C. albicans*). In both species, glycoside hydrolase treatment resulted in a higher viable dispersed population relative to treatment with antimicrobials alone (Fig 7A, Fig 7D). This was particularly notable for *C. albicans* biofilms, in which no viable fungi were detected in the dispersed population when treated with amphotericin B alone. In the case of combination treated *P. aeruginosa,* this increase in the planktonic population was countered by a decrease in the viable biofilm population (Fig 7B). This led to a 3 log_10_ reduction in viability of the total population being reached at 128 µg/mL and 256 µg/mL gentamicin when combined with glycoside hydrolase treatment, which was not achieved with the same concentrations of gentamicin alone (Fig 7C). Glycoside hydrolase and amphotericin B resulted in minor decreases in the viability of *C. albicans* biofilm populations (Fig 7E). This, taken alongside the increase in planktonic *C. albicans* when exposed to glycoside hydrolase meant that none of the tested treatments achieved biofilm eradication of the total population (Fig 7F). Treatment of *K. pneumoniae* biofilms with colistin and glycoside hydrolases was not explored, because *K. pneumoniae* biofilms were not dispersed by glycoside hydrolases (Fig 5B).

**Figure 7:**
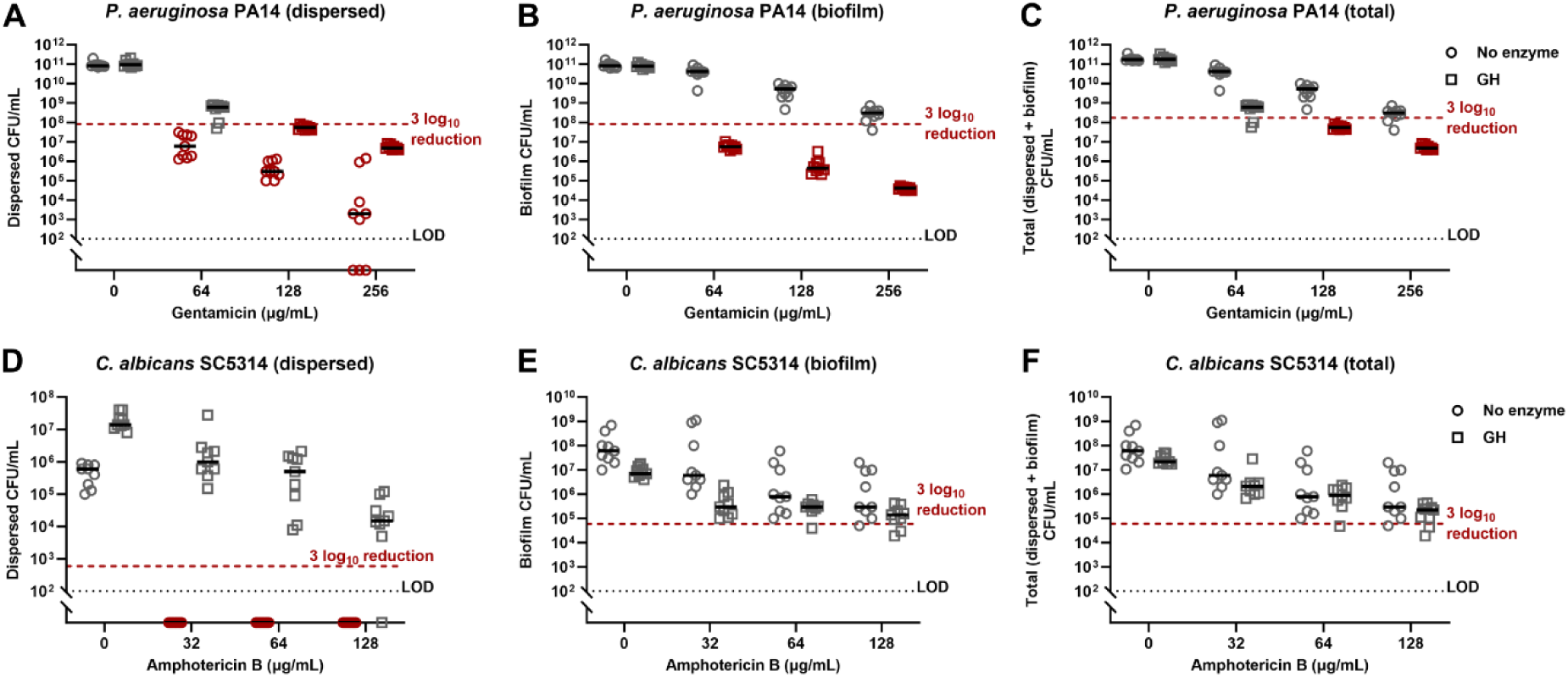
Treating *P. aeruginosa* and *C. albicans* with a combination of glycoside hydrolases and antimicrobials. Biofilms of *P. aeruginosa* or *C. albicans* biofilms were grown on segments of serum-coated endotracheal tube in SVAM for 48 hours. Endotracheal tubes were then transferred to fresh medium and treated with either 10% glycoside hydrolases (GH) and differing concentrations of either gentamicin (*P. aeruginosa*) or amphotericin B (*C. albicans*) for a further 24 hours. Viability of *P. aeruginosa* was determined by acquiring the endpoint CFUs of the dispersed **(A)**, biofilm **(B)**, which added together provide total viability of *P. aeruginosa* **(C).** Viability of *C. albicans* was determined by acquiring the endpoint CFUs of the dispersed **(D)**, biofilm **(E)**, which added together provide total viability of *C. albicans* **(F).** Red dotted line denotes the 3 log10 reduction threshold. Black dotted line denotes the limit of detection (LOD), n = 3 biological repeats, with 3 technical repeats for each biological replicate.

Proteinase K in combination with antimicrobials was the most effective treatment for both dispersing and eradicating biofilms of all species. The enzyme was most potent against *K. pneumoniae* when combined with colistin. This combination achieved almost complete eradication of dispersed (Fig 8D), biofilm (Fig 8E), and the total population (Fig 8F) in the endotracheal tube model. Thus, the combination of 64 µg/mL colistin and proteinase K was far superior to even 256 µg/mL colistin alone. Combining proteinase K with gentamicin enhanced clearance of both dispersed (Fig 8A) and biofilm associated (Fig 8B) *P. aeruginosa* compared to gentamicin alone. As with both DNase and glycoside hydrolase combination treatment, eradication of total *P. aeruginosa* was achieved by combining 128 µg/mL - 256 µg/mL gentamicin with proteinase K (Fig 8C). As observed with glycoside hydrolase combination treatment, proteinase K treatment increased the recovery of dispersed *C. albicans* (Fig 8G). However, unlike glycoside hydrolase combination treatment, proteinase K combined with amphotericin B led to a 3 log_10_ reduction in *C. albicans* biofilm viability relative to treating with amphotericin B alone (Fig 8H). We observed that 128 µg/mL amphotericin B in conjunction with proteinase K eradicated both the dispersed and biofilm *C. albicans* populations (Fig 8I).

**Figure 8:**
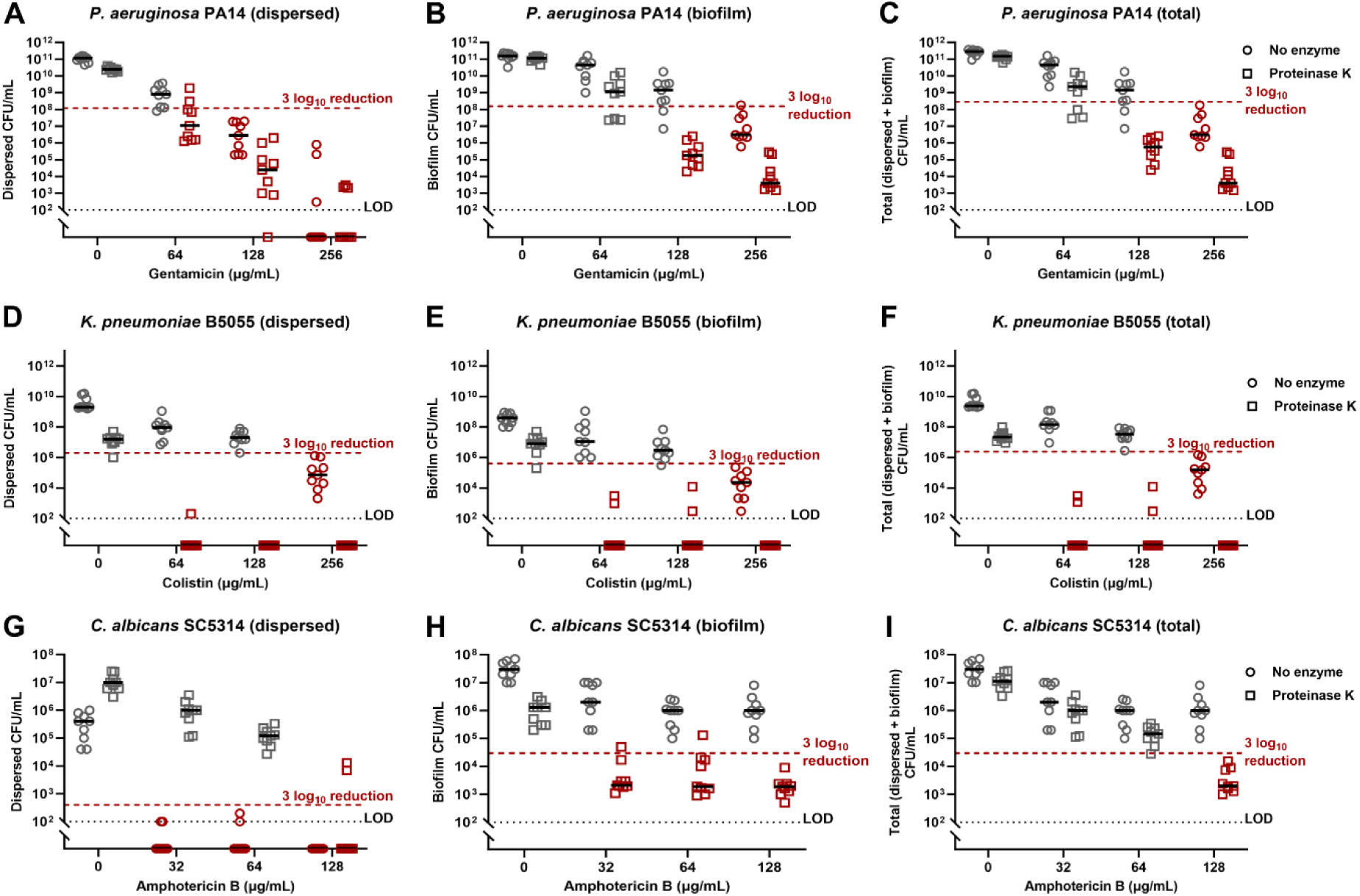
Treating biofilms grown in the *in vitro* endotracheal tube biofilm model with a combination of proteinase K and clinically relevant antimicrobials. *P. aeruginosa*, *K. pneumoniae*, or *C. albicans* biofilms were grown on segments of serum-coated endotracheal tube in SVAM for 48 hours. Endotracheal tubes were then transferred to fresh medium and treated with either 1 mg/mL proteinase K and differing concentrations of either gentamicin (*P.* aeruginosa), colistin (*K. pneumoniae*) or amphotericin B (*C. albicans*) for a further 24 hours. Viability of *P. aeruginosa* was determined by acquiring the endpoint CFUs of the dispersed cells **(A)** and remaining biofilm **(B)**, which added together provide total viability of *P. aeruginosa* (C). Viability of *K. pneumoniae* (D-F) and *C. albicans* (G-I), was similarly assessed. Red dotted line denotes the 3 log10 reduction threshold. Black dotted line denotes the limit of detection (LOD), n = 3 biological repeats, with 3 technical repeats for each biological replicate.

## Discussion

Ventilated associated pneumonia is broadly defined as a pneumonia occurring after more than 48 hours of mechanical ventilation. Ventilator-associated pneumonia mortality rates can reach 45% (58, 59). However, widespread differences in ventilator-associated pneumonia definition, diagnosis, and sampling account for large discrepancies in ventilator-associated pneumonia incidence and mortality rates globally (22, 58–62). The dangers of ventilator-associated pneumonia have been highlighted over recent years due to the coronavirus disease 2019 (COVID-19) pandemic, in which many hospitalised COVID-19 patients required long term mechanical ventilation (24). The reported incidence of ventilator-associated pneumonia in COVID-19 patients is much higher (50-80%) (63) than in non-COVID-19 patients (5-40%) (22, 64). Despite the need to study ventilator-associated pneumonia aetiology being higher than ever, the extensive development of host-mimicking media and *in vitro* and *ex vivo* infection models seen for conditions such as cystic fibrosis (34–41) has not transpired for ventilator-associated pneumonia, restricting ventilator-associated pneumonia microbiology research, and ultimately limiting our ability to combat the endotracheal tube biofilms that promote this disease.

Here, we present a novel *in vitro* endotracheal tube biofilm model, consisting of serum-coated endotracheal tube sections and a newly formulated growth medium to simulate the airway mucus of mechanically ventilated patients. We used this model to gain insights into biofilm structure of three important ventilator-associated pneumonia pathogens, and demonstrated that the model can be used to screen candidate therapies to combat endotracheal tube biofilms.

Endotracheal tubes are readily available for laboratory use, and serum-coating them ensured that the substrate in our model simulated the conditioning layer that coats endotracheal tubes following intubation (65). This layer is important for providing initial anchorage points to which microbes adhere. In medical device biofilms this layer consists of proteins such as fibronectin and laminin, and polysaccharides (66), all of which are present within FBS (67). Our SVAM medium was designed following extensive review of published literature (55). It contains many components absent from standard laboratory growth medium such as mucin, DNA, metals, GlcNAc, albumin and lysozyme which provide nutrition or alter the gene expression of microbial pathogens in the pulmonary inflammatory disease environment (68–75), and induce antimicrobial tolerance (70, 76–80).

The biofilms established by ventilator-associated pathogens were more difficult to eradicate when grown in our model than in a Calgary device, even if SVAM was the growth medium in the Calgary device. This mirrors previous findings: biofilms grown in our *ex vivo* cystic fibrosis biofilm model were more tolerant to colistin treatment than biofilms grown in cystic fibrosis-mimicking medium in a Calgary device (2). This increase in tolerance could be attributed to the serum-coated endotracheal tube upon which biofilms formed, as serum itself can induce antimicrobial tolerance (81).

From cryo-SEM of endotracheal tube biofilms, it was apparent that the model environment yielded very thick biofilm structures coated in thick matrix. This is consistent with *in vivo* data in other studies that have found up to 84% of endotracheal tubes extubated from patients were completely covered in confluent biofilm (82), and that this high coverage of biofilm increased the likelihood of ventilator-associated pneumonia (23). The microcolony structures observed in our biofilms are similar to those formed in sputum due to the presence of mucin and DNA (83), and the overall structure of the *P. aeruginosa* biofilms in our model – punctuated with gaps, giving a lacy or spongy appearance - resembled that seen in a cystic fibrosis lung biofilm model using light microscopy (41, 84, 85). For LB-grown *K. pneumoniae*, only small aggregates of biofilm could be found on endotracheal tubes, whilst SVAM resulted in extensive, matrix covered biofilms being observed. The rough cell surface of *K. pneumoniae* seen in SVAM is consistent with previous literature showing capsular polysaccharide on the cell surface of hypermucoviscous strains (86), such as *K. pneumoniae* B5055. The increased capsular polysaccharide in SVAM grown biofilms might be ascribed to the presence of glucose and iron in the SVAM medium, both of which are known to impact capsular polysaccharide biosynthesis in *K. pneumoniae* (87, 88).

The fungal pathogen *C. albicans* also displayed obvious morphological differences when grown in different endotracheal tube culture environments, favouring hyphal formation in YPD whilst remaining in yeast form or forming pseudohyphae in SVAM. GlcNAc, a component of SVAM, is known to induce filamentation of *C. albicans* (89). Serum, which coats the endotracheal tubes for both the YPD and SVAM conditions is also known to induce hyphal formation (90). However, the reason this may not occur in SVAM could be due to the presence of high concentrations of mucin, which can suppress filamentation even in the presence of other hyphal inducing conditions (91). Hyphal formation is associated with increased virulence of *C. albicans* whilst the yeast form is generally more associated with benign commensalism (92). It would be interesting to understand if the distinct morphology and structure of endotracheal tube biofilms contributes to the rare association of *C. albicans* pneumonia in ventilated patients, despite its frequent isolation from endotracheal tube biofilms (17, 20, 23, 93).

Since the biofilm matrix produced by ventilator-associated pneumonia pathogens varied depending on growth environment, matrix degrading enzyme susceptibility was also seen to be altered. Matrix-degrading enzymes could be powerful adjuvants to increase the ability of antimicrobials to eradicate biofilms, if the correct enzymes are chosen to ensure both dispersal and killing (94, 95). For example, DNase + gentamicin was tested for its ability to clear *P. aeruginosa* biofilms as rhDNase is commonly prescribed for mucus clearance in the CF airway, an environment in which *P. aeruginosa* biofilms are a common affliction (56, 57). Alone, DNase did not significantly disperse *P. aeruginosa* biofilms as judged by crystal violet assays; however DNase treatment did impact *P. aeruginosa* biofilms as it was seen to potentiate gentamicin killing of *P. aeruginosa* biofilms grown in the endotracheal tube model. Similar results of synergy have been reported for killing *P. aeruginosa* in cystic fibrosis sputum (96).

The glycoside hydrolase cocktail used for dispersal in this study consisted of α-amylase and cellulase, which hydrolyse α-1,4-glycosidic bonds and β-1,4-glycosidic bonds, respectively. These enzymes in conjunction with gentamicin have previously eradicated *P. aeruginosa* biofilms from murine chronic wounds (97), and this combination was also successful in our endotracheal tube model. Whilst this glycoside hydrolase cocktail significantly dispersed *C. albicans* biofilms grown in the model, it did not increase biofilm eradication when used in combination with amphotericin B relative to amphotericin B alone. In future, use of β-1,3-glucanase to degrade β-1,3-glucan, one of the most important matrix polysaccharides in *C. albicans* biofilms, may be a more appropriate candidate for testing (98, 99). Similarly, trials could be established with the use of bacteriophage-derived hydrolases that degrade the capsular polysaccharide of *K. pneumoniae* strains such as B5055 (100, 101).

In our model, we assessed viability of both the biofilm and dispersed populations after treatment with enzymes plus antimicrobials, as it is important to ensure that any biofilm dispersed from the endotracheal tube was sufficiently killed. Enzymatic degradation of biofilms without sufficient killing of dispersed cells could cause dissemination of infection (102). This is very important in the context of ventilator-associated pneumonia, which results from dispersal of biofilm deeper into the airways (16). Tightly controlled dosage and administration of matrix degrading enzymes and antimicrobial combinations, alongside the already routine regular suctioning of ventilated airways to clear mucus and dispersed pathogens (103), would be necessary if combination treatment were to be tried clinically.

In conclusion, through the development of SVAM growth medium combined with serum-coated endotracheal tube sections, we have produced an *in vitro* model that mimics the material, chemical, and nutritional environment surrounding ventilator-associated pathogens. As seen *in vivo*, we observed elevated antimicrobial tolerance and dense biofilm formation. As proof of concept that the model can be used to screen novel therapies, we assessed the ability of selected matrix-degrading enzymes to enhance the antibiofilm activity of clinically relevant antimicrobials. Some combinations, particularly proteinase K and colistin, were effective at eradicating both dispersed and endotracheal tube biofilm populations from the ventilator-associated environment. In future, this *in vitro* endotracheal tube (IVETT) model can be used to conduct more detailed explorations of ventilator-associated pathogen biology, to identify novel drug targets, and to screen candidate drugs or antimicrobial endotracheal tube coatings for likely efficacy before proceeding into *in vivo* trials.

## Acknowledgements

We would like to thank Jennifer Bevan and Matthew Pledge for their work aiding the development of the initial SVAM growth medium. We would also like to thank Claudia Simm for her advice and review of the manuscript. This work was funded by the University of Warwick and Monash University via the Monash-Warwick Alliance Training Programme in Emerging Superbug Threats, and by a University of Warwick School of Life Sciences Pump Priming Fund awarded to DW. AT was supported by a Future Fellowship from the Australian Research Council (FT190100733). TL acknowledges support of an Investigator Grant (2016330) from the Australian NHMRC. The authors wish to acknowledge the funding from the ESPRC, Grant number (EP/S021434/1) for the funding of the HR-CAT-SEM/Cryo SEM and to the nmRC for access to the facilities.

## Author contributions

The endotracheal tube model was originally conceived by FH, and developed by DW and FH. Experiments and hypotheses were developed by DW, FH, TL, AT and SB. DW and CP conducted experiments. DW analysed data. Funding was acquired by FH, TL, AT and DW. Project management was conducted by FH and DW.

## Supplementary methods

**Table S1:** Concentrations of each SVAM component in both their stocks and in the final SVAM medium.

| Component | Amount added to 500 mL SVAM base (3x strength) | Concentration in SVAM base (3x stock) | Concentration in final SVAM |
| --- | --- | --- | --- |
| CaCl<sub>2</sub> | 0.236 g | 4.26 mM | 1.42 mM |
| **Bovine serum albumin** | 10.5 g | 21 mg/mL | 7 mg/mL |
| **NaCl** | 3.33 g | 114 mM | 38 mM |
| **Na<sub>2</sub>HPO<sub>4</sub>** | 0.86 g | 12.12 mM | 4.04 mM |
| **NaH<sub>2</sub>PO<sub>4</sub>** | 0.206 g | 3.45 mM | 1.15 mM |
| **K<sub>2</sub>SO<sub>4</sub>** | 1.087 g | 12.48 mM | 4.16 mM |
| **KCl** | 0.131 g | 3.54 mM | 1.18 mM |
| **Casamino acids** | 7.5 g | 15 mg/mL | 5 mg/mL |
| **MOPS** | 3.139 g | 30 mM | 10 mM |
| **MgCl<sub>2</sub>·6H<sub>2</sub>O** | 0.725 g | 7.14 mM | 2.38 mM |
| **Glucose** | 1.08 g | 12 mM | 4 mM |
| **DPTA** | 0.009 g | 45 µM | 15 µM |
| **GlcNAc** | 5.25 g | 10.5 mg/mL | 3.5 mg/mL |
| **Fe(II)SO<sub>4</sub>·7H<sub>2</sub>O** | 1.5 mL of 17 mM stock | 51 µM | 17 µM |
| **ZnSO<sub>4</sub>·7H<sub>2</sub>O** | 1.5 mL of 95 mM stock | 285 µM | 95 µM |
| **CuSO<sub>4</sub>·5H<sub>2</sub>O** | 1.5 mL of 9.5 mM stock | 28.5 µM | 9.5 µM |
| **Sialic acid** | 1.5 mL of 4 mg/mL stock | 12 µg/mL | 4 µg/mL |
| **Component** | **Amount added to 80 mL (2x strength)** | **Concentration in 2x stock** | **Concentration in final SVAM** |
| **Porcine stomach mucin (Type III)** | 1.6 g | 20 mg/mL | 10 mg/mL |
| **Component** | **Amount added to 250 mL (10x strength)** | **Concentration in 10x stock** | **Concentration in final SVAM** |
| **Salmon sperm DNA salt** | 2.4 g | 9.6 mg/mL | 0.96 mg/mL |

**Table S2:** Suppliers and further information for SVAM components and enzymes used.

| **Component** | **Supplier** | **Further details** |
| --- | --- | --- |
| **CaCl<sub>2</sub>** | Alfa Aesar (L13191) |  |
| **Bovine serum albumin** | Sigma-Aldrich (A2153) |  |
| **NaCl** | Fisher Scientific (S/3160/60) |  |
| **Na<sub>2</sub>HPO<sub>4</sub>** | Acros organics (42437500) |  |
| **NaH<sub>2</sub>PO<sub>4</sub>** | Acros organics (389872500) |  |
| **K<sub>2</sub>SO<sub>4</sub>** | Acros organics (424222500) |  |
| **KCl** | Fisher Scientific (P/4280/53) |  |
| **Casamino acids** | VWR (J851) |  |
| **MOPS** | Acros organics (172635000) |  |
| **MgCl<sub>2</sub>·6H<sub>2</sub>O** | Sigma-Aldrich (M2670) |  |
| **Glucose** | Fisher Scientific (G/0500/53) |  |
| **DPTA** | Alfa Aesar (A10926) | Diethylenetriaminepentaacetic acid |
| **GlcNAc** | Sigma-Aldrich (A3286) | N-acetyl-D-glucosamine |
| **Fe(II)SO<sub>4</sub>·7H<sub>2</sub>O** | Sigma-Aldrich (F7002) |  |
| **ZnSO<sub>4</sub>·7H<sub>2</sub>O** | Thermo Scientific (205980010) |  |
| **CuSO<sub>4</sub>·5H<sub>2</sub>O** | Fisher Scientific (C/8520/53) |  |
| **Sialic acid** | Larodan (56-1150-13) | N-Acetyl-Neuraminic acid |
| **Salmon sperm DNA salt** | 31149 | Low molecular weight |
| **Porcine stomach mucin (Type III)** | Sigma-Aldrich (M1778) | Type III, bound sialic acid 0.5-1.5%, partially purified powder |
| **Human lysozyme** | Sigma-Aldrich (L1667) | Recombinant, expressed in rice, ≥100,000 units/mg protein |
| **DNase I** | Sigma-Aldrich (DN25) | From bovine pancreas, ≥400 Kunitz units/mg protein |
| **Cellulase** | Sigma-Aldrich (22178) | From *Aspergillus niger*, ~0.8 U/mg |
| **α-Amylase** | Sigma-Aldrich (10070) | From *Bacillus* sp., ~50 U/mg |
| **Proteinase K** | Sigma-Aldrich (SRE0047) | From *Triticum album*, ≥30 units/mg protein |
| **Fetal bovine serum** | Gibco (A3160801) | Origin: Brazil |
| **Amphotericin B** | Sigma-Aldrich (PHR1662) |  |
| **Gentamicin sulfate salt** | Sigma-Aldrich (G1264) |  |
| **Colistin sulfate salt** | Thermo Scientific (455390010) |  |

### Variants of SVAM medium

To investigate the effect of different SVAM components, variants of SVAM medium were made by supplementing a SVAM minimal medium stock with individual components. Table S3 details to preparation of the SVAM minimal medium stock. SVAM minimal medium stock is then diluted 1:3 with dH2O and filter sterilised to make SVAM base, which forms the base for all SVAM variants. Table S4 describes how the SVAM base is supplemented with additional components to produce SVAM variants.

**Table S3:**
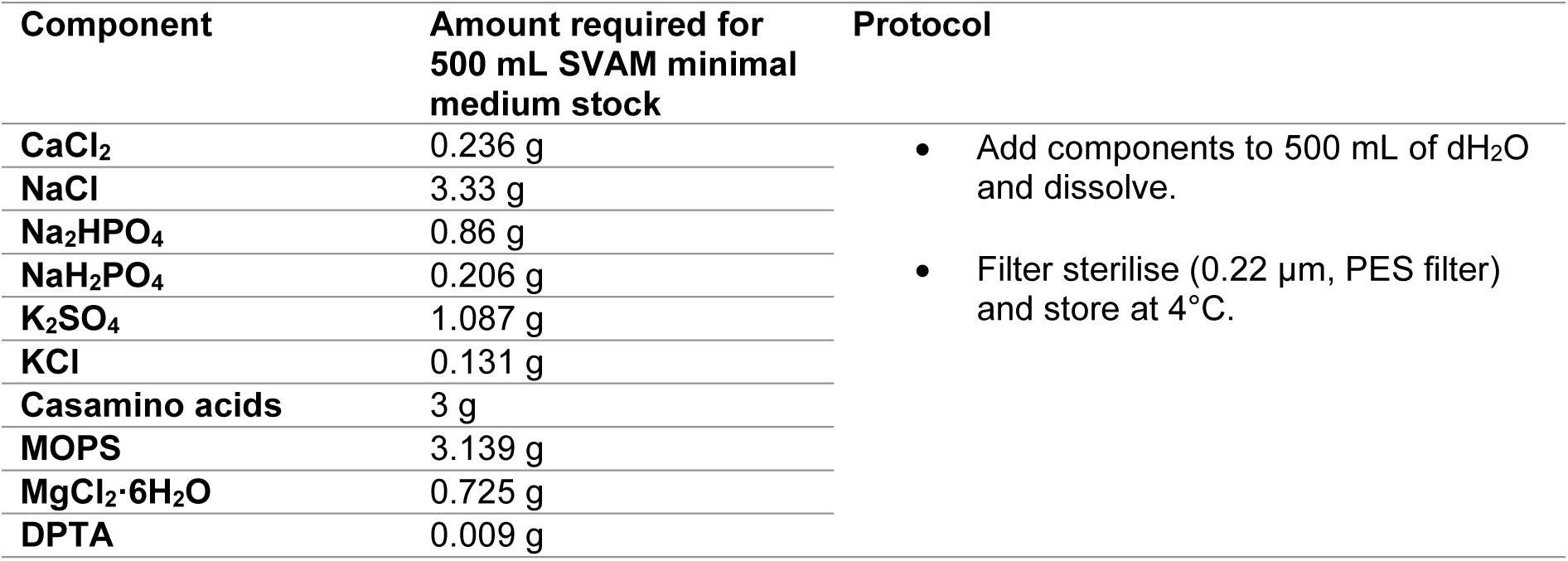
Protocol for preparation of SVAM minimal medium base.

**Table S4:** Protocol for preparation of SVAM medium variants.

| Component | Present in which SVAM variant (final concentration) | Supplementation protocol to make 500 mL SVAM variant. |
| --- | --- | --- |
| **CaCl<sub>2</sub>** | Base (1.42 mM) | SVAM minimal medium stock diluted 1:3 with dH<sub>2</sub>O to make SVAM base. |
| **NaCl** | Base (38 mM) |  |
| **Na<sub>2</sub>HPO<sub>4</sub>** | Base (4.04 mM) | Present in all variants as part of the SVAM minimal medium (base). |
| **NaH<sub>2</sub>PO<sub>4</sub>** | Base (1.15 mM) |  |
| **K<sub>2</sub>SO<sub>4</sub>** | Base (4.16 mM) |  |
| **KCl** | Base (1.18 mM) |  |
| **MOPS** | Base (10 mM) |  |
| **MgCl<sub>2</sub>·6H<sub>2</sub>O** | Base (2.38 mM) |  |
| **DPTA** | Base (15 µM) |  |
| **Casamino acids** | Base (2 mg/mL), casamino acids (5 mg/mL) | Part of base medium, a further 4.5 g added to 500 mL base to make casamino acid variant. |
| **Glucose** | Glucose (15 µM) | 1.08 g glucose dissolved in 500 mL base medium. |
| **GlcNAc** | GlcNAc (3.5 mg/mL) | 5.25 g GlcNAc dissolved in 500 mL base medium. |
| **Fe(II)SO<sub>4</sub>·7H<sub>2</sub>O** | Fe (17 µM) | 1.5 mL of 17 mM Fe(II)SO<sub>4</sub>·7H<sub>2</sub>O stock added to 500 mL base medium. |
| **ZnSO<sub>4</sub>·7H<sub>2</sub>O** | Zn (95 µM) | 1.5 mL of 9.5 mM CuSO<sub>4</sub>·5H<sub>2</sub>O stock added to 500 mL base medium. |
| **CuSO<sub>4</sub>·5H<sub>2</sub>O** | Cu (9.5 µM) | 1.5 mL of 95 mM ZnSO<sub>4</sub>·7H<sub>2</sub>O stock added to 500 mL base medium. |
| **Sialic acid** | Sialic acid (4 µg/mL) | 1.5 mL of 4 mg/mL sialic acid stock added to 500 mL base medium. |
| **Bovine serum albumin** | Albumin (7 mg/mL) | 10.5 g bovine serum albumin dissolved in 500 mL base medium. |
| **Porcine stomach mucin (Type III)** | Mucin (10 mg/mL) | 167 mL of SVAM minimal medium stock mixed with 250 mL of autoclaved 20 mg/mL PGM and 87 mL of dH<sub>2</sub>O. |
| **Salmon sperm DNA salt** | DNA (0.96 mg/mL) | 167 mL of SVAM minimal medium stock mixed with 50 mL of filter sterile 9.6 mg/mL salmon sperm DNA and 283 mL of dH<sub>2</sub>O. |

Following supplementation, all variants are pH adjusted to pH 6.7-6.8. As PGM cannot be filter sterilised, autoclaved PGM is added to filter sterilised SVAM minimal medium stock and dH_2_O for the mucin variant. All other variants are filter sterilised. The completed SVAM medium was also used in these experiments as a reference (SVAM), the medium for making SVAM medium is described in the Materials and Methods section of the research article.

### CFU/mL determination

Wells of 24 well plates were filled with 0.5 mL of LB (*P. aeruginosa, K. pneumoniae*), YPD (*C. albicans*), or SVAM medium variants, inoculated with 0.05 OD_600_ of either *P. aeruginosa*, *K. pneumoniae*, or *C. albicans*. Biofilms were then incubated for a further 24 hours at 37°C and 5% CO_2_. Following incubation, medium was removed and 1 mL of PBS added to wells. Biofilms were then sonicated for 15 minutes and then scraped with sterile pipette tips to remove biofilm. Cell suspensions were plated onto relevant growth medium and incubated for 24 hours at 37°C. CFU/mL was calculated to determine biofilm viability.

### Biofilm biomass determination

Wells of 24 well plates were filled with 0.5 mL of LB (*P. aeruginosa, K. pneumoniae*), YPD (*C. albicans*), or SVAM medium variants, inoculated with 0.05 OD_600_ of either *P. aeruginosa*, *K. pneumoniae*, or *C. albicans*. The 24 well plates were then incubated at 37°C, 5% CO_2_ for 48 hours to allow for biofilm formation. Following incubation, growth medium was removed and biofilms immersed in 0.5% (v/v) crystal violet for 15 minutes, washed in PBS and left to dry for 30 minutes. Stained biofilms were then immersed in 30% acetic acid for 15 minutes to solubilise crystal violet. Biomass was determined by reading absorbance at 550 nm.

**Figure S1:**
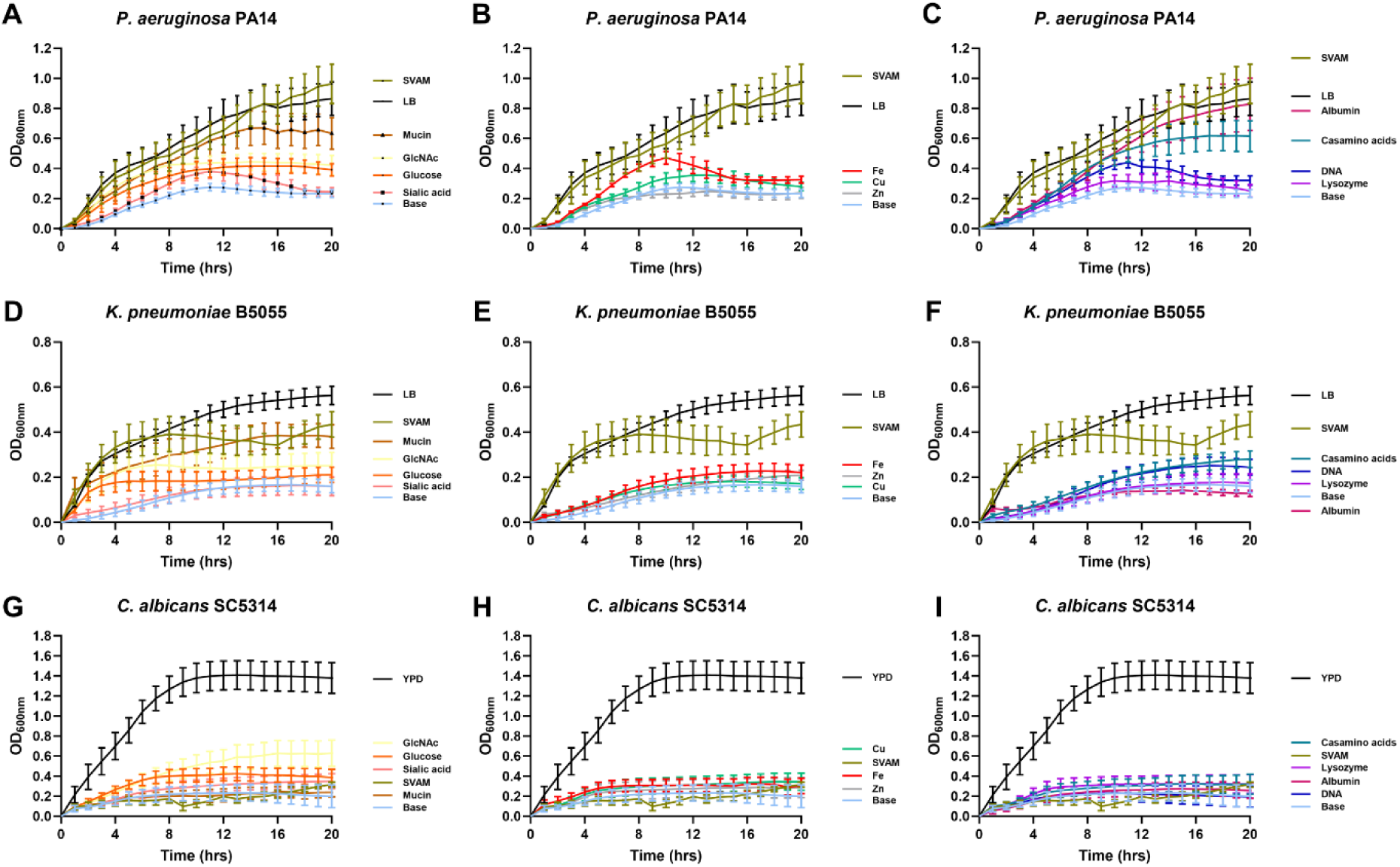
Effect of SVAM medium components on the growth of ventilator-associated pneumonia pathogens. *P. aeruginosa*, *K. pneumoniae*, and *C. albicans* were grown for 20 hours at 37°C. Growth curves plotting optical density at 600 nm (OD600nm) against time showing how mucin, GlcNAc, glucose and sialic acid affect the growth of *P. aeruginosa* **(A)**, *K. pneumoniae* **(D)**, and *C. albicans* **(G)**. Growth curves showing how iron (Fe), copper (Cu), and zinc (Zn) affect the growth of *P. aeruginosa* **(B)**, *K. pneumoniae* **(E)**, and *C. albicans* **(H).** Growth curves showing how casamino acids, DNA, lysozyme, and DNA affect the growth of *P. aeruginosa* **(C)**, *K. pneumoniae* **(F)**, and *C. albicans* **(I).** Standard laboratory medium (LB, *P. aeruginosa* and *K. pneumoniae*; YPD, *C. albicans*), SVAM, and minimal SVAM (base) were also shown on all growth curves for all species, n = 3 biological repeats, with 3 technical repeats for each biological replicate.

**Figure S2:**
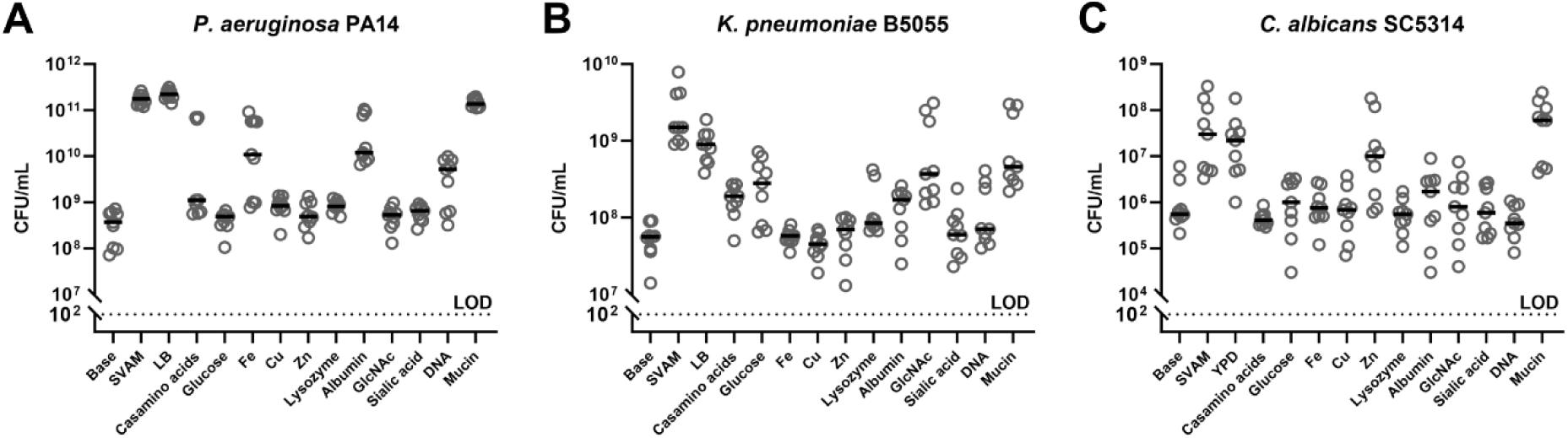
Characterising how components of SVAM medium affect the endpoint viability of ventilator-associated pneumonia pathogens. *P. aeruginosa*, *K. pneumoniae*, and *C. albicans* formed biofilms for 48 hours in either standard laboratory medium (LB or YPD), minimal SVAM (base), full SVAM (SVAM) or minimal SVAM supplemented with specific SVAM components for 48 hours. Endpoint CFUs of *P. aeruginosa* (A), *K. pneumoniae* (B), and *C. albicans* (C) were acquired to determine biofilm viability. LOD denotes limit of detection, n = 3 biological repeats, with 3 technical repeats for each biological replicate.

**Figure S3:**
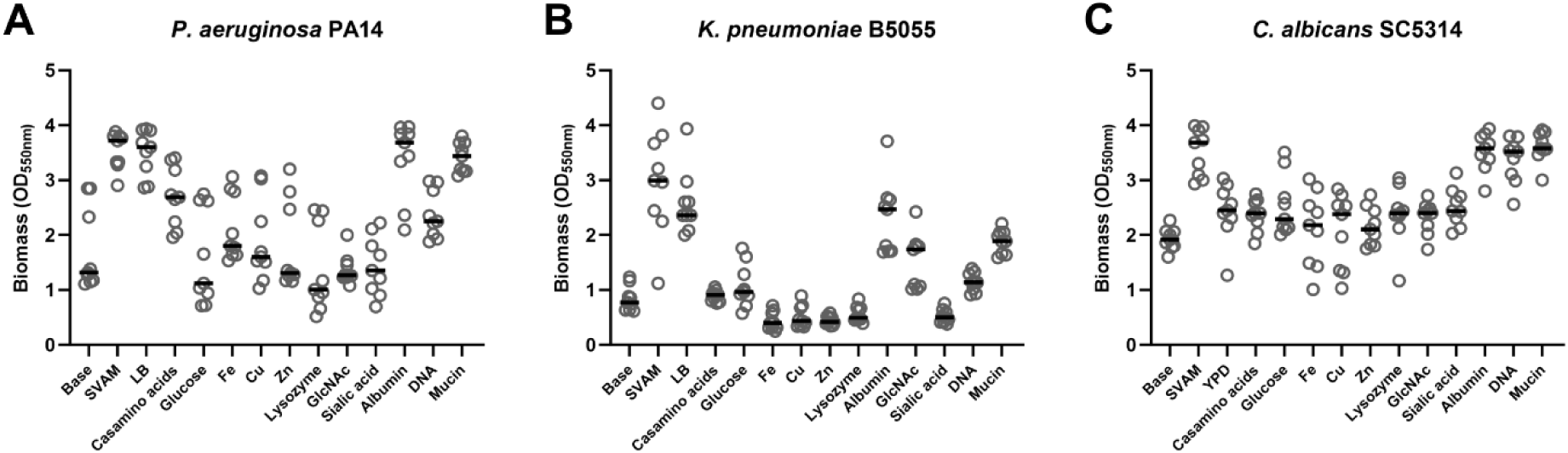
Characterising how components of SVAM medium affect the biomass of biofilms formed by ventilator-associated pneumonia pathogens. *P. aeruginosa*, *K. pneumoniae*, and *C. albicans* formed biofilms for 48 hours in either standard laboratory medium (LB or YPD), minimal SVAM (base), full SVAM (SVAM) or minimal SVAM supplemented with specific SVAM components for 48 hours. Crystal violet staining of *P. aeruginosa* (A), *K. pneumoniae* (B), and *C. albicans* (C) allowed for determination of biofilm biomass by spectral absorption of crystal violet at 550 nm (OD550nm). LOD denotes limit of detection, n = 3 biological repeats, with 3 technical repeats for each biological replicate.

## References

1. Donlan RM. Biofilms: microbial life on surfaces. Emerg Infect Dis. 2002;8(9):881–90.

2. Sweeney E, Sabnis A, Edwards AM, Harrison F. Effect of host-mimicking medium and biofilm growth on the ability of colistin to kill Pseudomonas aeruginosa. Microbiology (Reading). 2020;166(12):1171–80.

3. Anderl JN, Franklin MJ, Stewart PS. Role of antibiotic penetration limitation in Klebsiella pneumoniae biofilm resistance to ampicillin and ciprofloxacin. Antimicrob Agents Chemother. 2000;44(7):1818–24.

4. Sharma D, Misba L, Khan AU. Antibiotics versus biofilm: an emerging battleground in microbial communities. Antimicrob Resist Infect Control. 2019;8:76.

5. Gordon CA, Hodges, N. A., Marriott, C. Antibiotic interaction and diffusion through alginate and exopolysaccharide of cystic fibrosis derived Pseudomonas aeruginosa. J Antimicrob Chemother. 1988;22.

6. Nichols WW, Dorrington SM, Slack MP, Walmsley HL. Inhibition of tobramycin diffusion by binding to alginate. Antimicrob Agents Chemother. 1988;32(4):518–23.

7. Wood TK, Knabel SJ, Kwan BW. Bacterial persister cell formation and dormancy. Appl Environ Microbiol. 2013;79(23):7116–21.

8. Uruén C, Chopo-Escuin, G., Tommassen, J., Mainar-Jaime, R. C., Arenas, J. Biofilms as Promoters of Bacterial Antibiotic Resistance and Tolerance. Antibiotics (Basel). 2021;10(1).

9. Conibear TC, Collins SL, Webb JS. Role of mutation in Pseudomonas aeruginosa biofilm development. PLoS One. 2009;4(7):e6289.

10. Usui M, Yoshii Y, Thiriet-Rupert S, Ghigo JM, Beloin C. Intermittent antibiotic treatment of bacterial biofilms favors the rapid evolution of resistance. Commun Biol. 2023;6(1):275.

11. Magill SS, Edwards JR, Bamberg W, Beldavs ZG, Dumyati G, Kainer MA, Lynfield R, Maloney M, McAllister-Hollod L, Nadle J, Ray SM, Thompson DL, Wilson LE, Fridkin SK, Emerging Infections Program Healthcare-Associated I, Antimicrobial Use Prevalence Survey T. Multistate point-prevalence survey of health care-associated infections. N Engl J Med. 2014;370(13):1198–208.

12. Ferreira Tde O, Koto RY, Leite GF, Klautau GB, Nigro S, Silva CB, Souza AP, Mimica MJ, Cesar RG, Salles MJ. Microbial investigation of biofilms recovered from endotracheal tubes using sonication in intensive care unit pediatric patients. Braz J Infect Dis. 2016;20(5):468–75.

13. American Thoracic S, Infectious Diseases Society of A. Guidelines for the management of adults with hospital-acquired, ventilator-associated, and healthcare-associated pneumonia. Am J Respir Crit Care Med. 2005;171(4):388–416.

14. Durairaj L, Mohamad Z, Launspach JL, Ashare A, Choi JY, Rajagopal S, Doern GV, Zabner J. Patterns and density of early tracheal colonization in intensive care unit patients. J Crit Care. 2009;24(1):114–21.

15. Pneumatikos IA, Dragoumanis CK, Bouros DE. Ventilator-associated pneumonia or endotracheal tube-associated pneumonia? An approach to the pathogenesis and preventive strategies emphasizing the importance of endotracheal tube. Anesthesiology. 2009;110(3):673–80.

16. Diaconu O, Siriopol I, Polosanu LI, Grigoras I. Endotracheal Tube Biofilm and its Impact on the Pathogenesis of Ventilator-Associated Pneumonia. Journal of critical care medicine (Universitatea de Medicina si Farmacie din Targu-Mures). 2018;4(2):50–5.

17. Hotterbeekx A, Xavier BB, Bielen K, Lammens C, Moons P, Schepens T, Ieven M, Jorens PG, Goossens H, Kumar-Singh S, Malhotra-Kumar S. The endotracheal tube microbiome associated with Pseudomonas aeruginosa or Staphylococcus epidermidis. Sci Rep. 2016;6:36507.

18. Cifuentes EA, Sierra MA, Yepes AF, Baldion AM, Rojas JA, Alvarez-Moreno CA, Anzola JM, Zambrano MM, Huertas MG. Endotracheal tube microbiome in hospitalized patients defined largely by hospital environment. Respir Res. 2022;23(1):168.

19. Adair CG, Gorman SP, Feron BM, Byers LM, Jones DS, Goldsmith CE, Moore JE, Kerr JR, Curran MD, Hogg G, Webb CH, McCarthy GJ, Milligan KR. Implications of endotracheal tube biofilm for ventilator-associated pneumonia. Intensive Care Med. 1999;25(10):1072–6.

20. Zakharkina T, Martin-Loeches I, Matamoros S, Povoa P, Torres A, Kastelijn JB, Hofstra JJ, de Wever B, de Jong M, Schultz MJ, Sterk PJ, Artigas A, Bos LDJ. The dynamics of the pulmonary microbiome during mechanical ventilation in the intensive care unit and the association with occurrence of pneumonia. Thorax. 2017;72(9):803–10.

21. Arayasukawat P, So-Ngern, A., Reechaipichitkul, W., Chumpangern, W., Arunsurat, I., Ratanawatkul, P., Chuennok, W. Microorganisms and clinical outcomes of early-and late-onset ventilator-associated pneumonia at Srinagarind Hospital, a tertiary center in Northeastern Thailand. BMC Pulm Med. 2021;21(1).

22. Alves D, Grainha T, Pereira MO, Lopes SP. Antimicrobial materials for endotracheal tubes: A review on the last two decades of technological progress. Acta Biomater. 2023;158:32–55.

23. Thorarinsdottir HR, Kander T, Holmberg A, Petronis S, Klarin B. Biofilm formation on three different endotracheal tubes: a prospective clinical trial. Crit Care. 2020;24(1):382.

24. van Duijnhoven M, Fleuren-Janssen M, van Osch F, LeNoble J. A Predominant Cause of Recurrence of Ventilator-Associated Pneumonia in Patients with COVID-19 Are Relapses. J Clin Med. 2023;12(18).

25. Marcut L, Manescu Paltanea V, Antoniac A, Paltanea G, Robu A, Mohan AG, Grosu E, Corneschi I, Bodog AD. Antimicrobial Solutions for Endotracheal Tubes in Prevention of Ventilator-Associated Pneumonia. Materials (Basel). 2023;16(14).

26. Piraino T. The Role of Specialty Tubes in Preventing Ventilator-Associated Events. Respir Care. 2019;64(8):980–5.

27. Ersoy SC, Heithoff DM, Barnes Lt, Tripp GK, House JK, Marth JD, Smith JW, Mahan MJ. Correcting a Fundamental Flaw in the Paradigm for Antimicrobial Susceptibility Testing. EBioMedicine. 2017;20:173–81.

28. Tetz G, Tetz V. Overcoming Antibiotic Resistance with Novel Paradigms of Antibiotic Selection. Microorganisms. 2022;10(12).

29. Tetz GV, Kardava KM, Vecherkovskaya MF, Tsifansky MD, Tetz VV. Treatment of chronic relapsing urinary tract infection with antibiotics selected by AtbFinder. Urol Case Rep. 2023;46:102312.

30. Thieme L, Hartung, A., Tramm, K., Klinger-Strobel, M., Jandt, K. D., Makarewicz, O., Pletz, M. W. MBEC Versus MBIC: the Lack of Differentiation between Biofilm Reducing and Inhibitory Effects as a Current Problem in Biofilm Methodology. Biol Proced Online. 2019;21(18).

31. Moskowitz SM, Foster JM, Emerson J, Burns JL. Clinically feasible biofilm susceptibility assay for isolates of Pseudomonas aeruginosa from patients with cystic fibrosis. J Clin Microbiol. 2004;42(5):1915–22.

32. Ceri H, Olson ME, Stremick C, Read RR, Morck D, Buret A. The Calgary Biofilm Device: new technology for rapid determination of antibiotic susceptibilities of bacterial biofilms. J Clin Microbiol. 1999;37(6):1771–6.

33. Smith S, Waters V, Jahnke N, Ratjen F. Standard versus biofilm antimicrobial susceptibility testing to guide antibiotic therapy in cystic fibrosis. Cochrane Database Syst Rev. 2020;6(6):CD009528.

34. Ghani M, Soothill JS. Ceftazidime, gentamicin, and rifampicin, in combination, kill biofilms of mucoid Pseudomonas aeruginosa. Can J Microbiol. 1997;43:999–1004.

35. Sriramulu DD, Lunsdorf H, Lam JS, Romling U. Microcolony formation: a novel biofilm model of Pseudomonas aeruginosa for the cystic fibrosis lung. J Med Microbiol. 2005;54(Pt 7):667–76.

36. Fung C, Naughton S, Turnbull L, Tingpej P, Rose B, Arthur J, Hu H, Harmer C, Harbour C, Hassett DJ, Whitchurch CB, Manos J. Gene expression of Pseudomonas aeruginosa in a mucin-containing synthetic growth medium mimicking cystic fibrosis lung sputum. J Med Microbiol. 2010;59(Pt 9):1089–100.

37. Palmer KL, Aye LM, Whiteley M. Nutritional cues control Pseudomonas aeruginosa multicellular behavior in cystic fibrosis sputum. J Bacteriol. 2007;189(22):8079–87.

38. Turner KH, Wessel AK, Palmer GC, Murray JL, Whiteley M. Essential genome of Pseudomonas aeruginosa in cystic fibrosis sputum. Proc Natl Acad Sci U S A. 2015;112(13):4110–5.

39. Ruhluel D, O’Brien S, Fothergill JL, Neill DR. Development of liquid culture media mimicking the conditions of sinuses and lungs in cystic fibrosis and health. F1000Research. 2022.

40. Harrison F, Diggle SP. An ex vivo lung model to study bronchioles infected with Pseudomonas aeruginosa biofilms. Microbiology (Reading). 2016;162(10):1755–60.

41. Harrington NE, Sweeney E, Harrison F. Building a better biofilm - Formation of in vivo-like biofilm structures by Pseudomonas aeruginosa in a porcine model of cystic fibrosis lung infection. Biofilm. 2020;2:100024.

42. Harrington NE, Sweeney E, Alav I, Allen F, Moat J, Harrison F. Antibiotic Efficacy Testing in an Ex vivo Model of Pseudomonas aeruginosa and Staphylococcus aureus Biofilms in the Cystic Fibrosis Lung. J Vis Exp. 2021(167).

43. Harrington NE, Littler JL, Harrison F. Transcriptome Analysis of Pseudomonas aeruginosa Biofilm Infection in an Ex Vivo Pig Model of the Cystic Fibrosis Lung. Appl Environ Microbiol. 2022;88(3):e0178921.

44. O’Toole GA, Crabbe A, Kummerli R, LiPuma JJ, Bomberger JM, Davies JC, Limoli D, Phelan VV, Bliska JB, DePas WH, Dietrich LE, Hampton TH, Hunter R, Khursigara CM, Price-Whelan A, Ashare A, Cramer RA, Goldberg JB, Harrison F, Hogan DA, Henson MA, Madden DR, Mayers JR, Nadell C, Newman D, Prince A, Rivett DW, Schwartzman JD, Schultz D, Sheppard DC, Smyth AR, Spero MA, Stanton BA, Turner PE, van der Gast C, Whelan FJ, Whitaker R, Whiteson K. Model Systems to Study the Chronic, Polymicrobial Infections in Cystic Fibrosis: Current Approaches and Exploring Future Directions. mBio. 2021;12(5):e0176321.

45. Luna CM, Sibila O, Agusti C, Torres A. Animal models of ventilator-associated pneumonia. Eur Respir J. 2009;33(1):182–8.

46. Pulido L, Burgos, D., García, J. M., Luna, C. M. Does animal model on ventilator-associated pneumonia reflect physiopathology of sepsis mechanisms in humans? Ann Transl Med. 2017;5(22).

47. Torres A, Niederman MS, Chastre J, Ewig S, Fernandez-Vandellos P, Hanberger H, Kollef M, Li Bassi G, Luna CM, Martin-Loeches I, Paiva JA, Read RC, Rigau D, Timsit JF, Welte T, Wunderink R. International ERS/ESICM/ESCMID/ALAT guidelines for the management of hospital-acquired pneumonia and ventilator-associated pneumonia: Guidelines for the management of hospital-acquired pneumonia (HAP)/ventilator-associated pneumonia (VAP) of the European Respiratory Society (ERS), European Society of Intensive Care Medicine (ESICM), European Society of Clinical Microbiology and Infectious Diseases (ESCMID) and Asociacion Latinoamericana del Torax (ALAT). Eur Respir J. 2017;50(3).

48. Boisson M, Mimoz O, Hadzic M, Marchand S, Adier C, Couet W, Gregoire N. Pharmacokinetics of intravenous and nebulized gentamicin in critically ill patients. J Antimicrob Chemother. 2018;73(10):2830–7.

49. Ramadan RA, Bedawy, A. M., Negm, E. M., Hassan, T. H., Ibrahim, D. A., ElSheikh, S. M., Amer, R. M.. Carbapenem-Resistant Klebsiella pneumoniae Among Patients with Ventilator-Associated Pneumonia: Evaluation of Antibiotic Combinations and Susceptibility to New Antibiotics. Infect Drug Resist. 2022;6(15).

50. Du H, Wei L, Li W, Huang B, Liu Y, Ye X, Zhang S, Wang T, Chen Y, Chen D, Liu J. Effect of Nebulized Amphotericin B in Critically ill Patients With Respiratory Candida spp. De-colonization: A Retrospective Analysis. Front Med (Lausanne). 2021;8:723904.

51. Henderson AG, Ehre C, Button B, Abdullah LH, Cai LH, Leigh MW, DeMaria GC, Matsui H, Donaldson SH, Davis CW, Sheehan JK, Boucher RC, Kesimer M. Cystic fibrosis airway secretions exhibit mucin hyperconcentration and increased osmotic pressure. J Clin Invest. 2014;124(7):3047–60.

52. Powell J, Garnett JP, Mather MW, Cooles FAH, Nelson A, Verdon B, Scott J, Jiwa K, Ruchaud-Sparagano MH, Cummings SP, Perry JD, Wright SE, Wilson JA, Pearson J, Ward C, Simpson AJ. Excess Mucin Impairs Subglottic Epithelial Host Defense in Mechanically Ventilated Patients. Am J Respir Crit Care Med. 2018;198(3):340–9.

53. Baker EH, Clark N, Brennan AL, Fisher DA, Gyi KM, Hodson ME, Philips BJ, Baines DL, Wood DM. Hyperglycemia and cystic fibrosis alter respiratory fluid glucose concentrations estimated by breath condensate analysis. J Appl Physiol (1985). 2007;102(5):1969-75.

54. Philips BJ, Redman J, Brennan A, Wood D, Holliman R, Baines D, Baker EH. Glucose in bronchial aspirates increases the risk of respiratory MRSA in intubated patients. Thorax. 2005;60(9):761–4.

55. Walsh D, Bevan J, Harrison F. How Does Airway Surface Liquid Composition Vary in Different Pulmonary Diseases, and How Can We Use This Knowledge to Model Microbial Infections? Preprints. 2024.

56. Suri R. The use of human deoxyribonuclease (rhDNase) in the management of cystic fibrosis. BioDrugs. 2005;19(3):135–44.

57. Pressler T. Review of recombinant human deoxyribonuclease (rhDNase) in the management of patients with cystic fibrosis. Biologics. 2008;2(4):611–7.

58. Mehta A, Bhagat R. Preventing Ventilator-Associated Infections. Clin Chest Med. 2016;37(4):683–92.

59. Nora D, Povoa P. Antibiotic consumption and ventilator-associated pneumonia rates, some parallelism but some discrepancies. Ann Transl Med. 2017;5(22):450.

60. Dudeck MA, Horan TC, Peterson KD, Allen-Bridson K, Morrell G, Anttila A, Pollock DA, Edwards JR. National Healthcare Safety Network report, data summary for 2011, device-associated module. Am J Infect Control. 2013;41(4):286–300.

61. Koulenti D, Tsigou E, Rello J. Nosocomial pneumonia in 27 ICUs in Europe: perspectives from the EU-VAP/CAP study. Eur J Clin Microbiol Infect Dis. 2017;36(11):1999–2006.

62. Rosenthal VD, Rodrigues C, Alvarez-Moreno C, Madani N, Mitrev Z, Ye G, Salomao R, Ulger F, Guanche-Garcell H, Kanj SS, Cuellar LE, Higuera F, Mapp T, Fernandez-Hidalgo R, members I. Effectiveness of a multidimensional approach for prevention of ventilator-associated pneumonia in adult intensive care units from 14 developing countries of four continents: findings of the International Nosocomial Infection Control Consortium. Crit Care Med. 2012;40(12):3121-8.

63. Gragueb-Chatti I, Hyvernat H, Leone M, Agard G, Peres N, Guervilly C, Boucekine M, Hamidi D, Papazian L, Dellamonica J, Lopez A, Hraiech S. Incidence, Outcomes and Risk Factors of Recurrent Ventilator Associated Pneumonia in COVID-19 Patients: A Retrospective Multicenter Study. J Clin Med. 2022;11(23).

64. Papazian L, Klompas M, Luyt CE. Ventilator-associated pneumonia in adults: a narrative review. Intensive Care Med. 2020;46(5):888–906.

65. Shein AMS, Wannigama DL, Higgins PG, Hurst C, Abe S, Hongsing P, Chantaravisoot N, Saethang T, Luk-In S, Liao T, Nilgate S, Rirerm U, Kueakulpattana N, Laowansiri M, Srisakul S, Muhummudaree N, Techawiwattanaboon T, Gan L, Xu C, Kupwiwat R, Phattharapornjaroen P, Rojanathanes R, Leelahavanichkul A, Chatsuwan T. Novel colistin-EDTA combination for successful eradication of colistin-resistant Klebsiella pneumoniae catheter-related biofilm infections. Sci Rep. 2021;11(1):21676.

66. Khatoon Z, McTiernan CD, Suuronen EJ, Mah TF, Alarcon EI. Bacterial biofilm formation on implantable devices and approaches to its treatment and prevention. Heliyon. 2018;4(12):e01067.

67. Lee DY, Lee SY, Yun SH, Jeong JW, Kim JH, Kim HW, Choi JS, Kim GD, Joo ST, Choi I, Hur SJ. Review of the Current Research on Fetal Bovine Serum and the Development of Cultured Meat. Food Sci Anim Resour. 2022;42(5):775–99.

68. Venkatakrishnan V, Packer NH, Thaysen-Andersen M. Host mucin glycosylation plays a role in bacterial adhesion in lungs of individuals with cystic fibrosis. Expert Rev Respir Med. 2013;7(5):553–76.

69. Lewenza S, Johnson L, Charron-Mazenod L, Hong M, Mulcahy-O’Grady H. Extracellular DNA controls expression of Pseudomonas aeruginosa genes involved in nutrient utilization, metal homeostasis, acid pH tolerance and virulence. J Med Microbiol. 2020;69(6):895–905.

70. Wilton M, Charron-Mazenod L, Moore R, Lewenza S. Extracellular DNA Acidifies Biofilms and Induces Aminoglycoside Resistance in Pseudomonas aeruginosa. Antimicrob Agents Chemother. 2016;60(1):544–53.

71. Feingold DS, Goldman JN, Kuritz HM. Locus of the action of serum and the role of lysozyme in the serum bactericidal reaction. J Bacteriol. 1968;96(6):2118–26.

72. Reid DW, Carroll V, O’May C, Champion A, Kirov SM. Increased airway iron as a potential factor in the persistence of Pseudomonas aeruginosa infection in cystic fibrosis. Eur Respir J. 2007;30:286–92.

73. Sunder-Plassmann G, Patruta SI, Horl WH. Pathobiology of the role of iron in infection. Am J Kidney Dis. 1999;34(4 Suppl 2):S25-9.

74. Egesten A, Frick IM, Mörgelin M, Olin AI, Björck L. Binding of Albumin Promotes Bacterial Survival at the Epithelial Surface. J Biol Chem. 2011;286(4):2469–76.

75. Kruczek C, Wachtel M, Alabady MS, Payton PR, Colmer-Hamood JA, Hamood AN. Serum albumin alters the expression of iron-controlled genes in Pseudomonas aeruginosa. Microbiology (Reading). 2012;158(Pt 2):353–67.

76. Sebbag L, Broadbent VL, Kenne DE, Perrin AL, Mochel JP. Albumin in Tears Modulates Bacterial Susceptibility to Topical Antibiotics in Ophthalmology. Front Med (Lausanne). 2021;8:663212.

77. Rodrigues AG, Araujo R, Pina-Vaz C. Human albumin promotes germination, hyphal growth and antifungal resistance by Aspergillus fumigatus. Med Mycol. 2005;43(8):711–7.

78. Lewenza S. Extracellular DNA-induced antimicrobial peptide resistance mechanisms in Pseudomonas aeruginosa. Front Microbiol. 2013;4:21.

79. Samad T, Co JY, Witten J, Ribbeck K. Mucus and Mucin Environments Reduce the Efficacy of Polymyxin and Fluoroquinolone Antibiotics against Pseudomonas aeruginosa. ACS Biomater Sci Eng. 2019;5(3):1189–94.

80. Landry RM, An D, Hupp JT, Singh PK, Parsek MR. Mucin-Pseudomonas aeruginosa interactions promote biofilm formation and antibiotic resistance. Mol Microbiol. 2006;59(1):142–51.

81. Morrison JM, Chojnacki M, Fadrowski JJ, Bauza C, Dunman PM, Dudas RA, Goldenberg NA, Berman DM. Serum-Associated Antibiotic Tolerance in Pediatric Clinical Isolates of Pseudomonas aeruginosa. J Pediatric Infect Dis Soc. 2020;9(6):671–9.

82. Sottile FD, Marrie TJ, Prough DS, Hobgood CD, Gower DJ, Webb LX, Costerton JW, Gristina AG. Nosocomial pulmonary infection: possible etiologic significance of bacterial adhesion to endotracheal tubes. Crit Care Med. 1986;14(4):265–70.

83. Guillaume O, Butnarasu, C., Visentin, S., Reimhult, E. Interplay between biofilm microenvironment and pathogenicity of Pseudomonas aeruginosa in cystic fibrosis lung chronic infection. Biofilm. 2022;4.

84. Jennings LK, Dreifus, J. E., Reichhardt, C., Storek, K. M., Secor, P. R., Wozniak, D. J., Hisert, K.B., Parsek, M. R.. Pseudomonas aeruginosa aggregates in cystic fibrosis sputum produce exopolysaccharides that likely impede current therapies. Cell Rep. 2021;23(34).

85. Roberts AEL, Powell LC, Pritchard MF, Thomas DW, Jenkins RE. Anti-pseudomonad Activity of Manuka Honey and Antibiotics in a Specialized ex vivo Model Simulating Cystic Fibrosis Lung Infection. Front Microbiol. 2019;10:869.

86. Cai R, Wang G, Le S, Wu M, Cheng M, Guo Z, Ji Y, Xi H, Zhao C, Wang X, Xue Y, Wang Z, Zhang H, Fu Y, Sun C, Feng X, Lei L, Yang Y, Ur Rahman S, Liu X, Han W, Gu J. Three Capsular Polysaccharide Synthesis-Related Glucosyltransferases, GT-1, GT-2 and WcaJ, Are Associated With Virulence and Phage Sensitivity of Klebsiella pneumoniae. Front Microbiol. 2019;10:1189.

87. Lin CT, Chen YC, Jinn TR, Wu CC, Hong YM, Wu WH. Role of the cAMP-dependent carbon catabolite repression in capsular polysaccharide biosynthesis in Klebsiella pneumoniae. PLoS One. 2013;8(2):e54430.

88. Zhu J, Wang T, Chen L, Du H. Virulence Factors in Hypervirulent Klebsiella pneumoniae. Front Microbiol. 2021;12:642484.

89. Naseem S, Gunasekera A, Araya E, Konopka JB. N-acetylglucosamine (GlcNAc) induction of hyphal morphogenesis and transcriptional responses in Candida albicans are not dependent on its metabolism. J Biol Chem. 2011;286(33):28671–80.

90. Feng Q, Summers, E., Guo, B., Fink, G. Ras signaling is required for serum-induced hyphal differentiation in Candida albicans. J Bacteriol. 1999;181(20).

91. Valle Arevalo A, Nobile CJ. Interactions of microorganisms with host mucins: a focus on Candida albicans. FEMS Microbiol Rev. 2020;44(5):645–54.

92. Noble SM, Gianetti, B. A., Witchley, J. N. Candida albicans cell type switches and functional plasticity in the mammalian host. Nat Rev Microbiol. 2017;15(2).

93. Maes M, Higginson E, Pereira-Dias J, Curran MD, Parmar S, Khokhar F, Cuchet-Lourenco D, Lux J, Sharma-Hajela S, Ravenhill B, Hamed I, Heales L, Mahroof R, Soderholm A, Forrest S, Sridhar S, Brown NM, Baker S, Navapurkar V, Dougan G, Scott JB, Morris AC. Correction to: Ventilator-associated pneumonia in critically ill patients with COVID-19. Crit Care. 2021;25(1):130.

94. Kovach KN, Fleming D, Wells MJ, Rumbaugh KP, Gordon VD. Specific Disruption of Established Pseudomonas aeruginosa Biofilms Using Polymer-Attacking Enzymes. Langmuir. 2020;36(6):1585–95.

95. Fleming D, Rumbaugh KP. Approaches to Dispersing Medical Biofilms. Microorganisms. 2017;5(2).

96. Alipour M, Suntres ZE, Omri A. Importance of DNase and alginate lyase for enhancing free and liposome encapsulated aminoglycoside activity against Pseudomonas aeruginosa. J Antimicrob Chemother. 2009;64(2):317–25.

97. Fleming D, Chahin L, Rumbaugh K. Glycoside Hydrolases Degrade Polymicrobial Bacterial Biofilms in Wounds. Antimicrob Agents Chemother. 2017;61(2).

98. Tan Y, Ma S, Ding T, Ludwig R, Lee J, Xu J. Enhancing the Antibiofilm Activity of beta-1,3-Glucanase-Functionalized Nanoparticles Loaded With Amphotericin B Against Candida albicans Biofilm. Front Microbiol. 2022;13:815091.

99. Nett J, Lincoln L, Marchillo K, Massey R, Holoyda K, Hoff B, VanHandel M, Andes D. Putative role of beta-1,3 glucans in Candida albicans biofilm resistance. Antimicrob Agents Chemother. 2007;51(2):510–20.

100. Dunstan RA, Bamert, R. S., Belousoff, M. J., Short, F. L., Barlow, C. K., Pickard, D. J., Wilksch, J. J., Schittenhelm, R. B., Strugnell, R. A., Dougan, G., Lithgow T. Mechanistic Insights into the Capsule-Targeting Depolymerase from a Klebsiella pneumoniae Bacteriophage. Microbiol Spectr. 2021;9(1).

101. Dunstan RA, Bamert RS, Tan KS, Imbulgoda U, Barlow CK, Taiaroa G, Pickard DJ, Schittenhelm RB, Dougan G, Short FL, Lithgow T. Epitopes in the capsular polysaccharide and the porin OmpK36 receptors are required for bacteriophage infection of Klebsiella pneumoniae. Cell Rep. 2023;42(6):112551.

102. Fleming D, Rumbaugh K. The Consequences of Biofilm Dispersal on the Host. Sci Rep. 2018;8(1):10738.

103. Ardehali SH, Fatemi A, Rezaei SF, Forouzanfar MM, Zolghadr Z. The Effects of Open and Closed Suction Methods on Occurrence of Ventilator Associated Pneumonia; a Comparative Study. Arch Acad Emerg Med. 2020;8(1):e8.

